# The *Caenorhabditis elegans* HAM-1 protein modifies G protein signaling and membrane extension to reverse the polarity of asymmetric cell division

**DOI:** 10.1101/504787

**Authors:** Jerome Teuliere, Gian Garriga

## Abstract

Asymmetric divisions often produce daughter cells that differ in both fate and size. The *Caenorhabditis elegans* HAM-1 protein regulates both daughter cell fate and daughter cell size asymmetry (DCSA) in a subset of asymmetric divisions. Here we focus on the divisions of the Q.a and Q.p neuroblasts, which use distinct mechanisms to divide with opposite polarity. Q.a divides by a *ham-1-*dependent, spindle-independent, myosin-dependent mechanism to produce a smaller anterior daughter that dies, whereas Q.p divides by a *ham-1*-independent, spindle-dependent, myosin-independent mechanism to produce a smaller posterior daughter that dies. Despite these differences, we found that membrane extension at the posterior of Q.a and at the anterior of Q.p promoted DCSA in these cells by a Wiscott-Aldrich protein (WASp)-dependent mechanism and that in *ham-1* mutant Q.a divisions, the polarity of this extension was reversed. In addition, the spindle moved posteriorly during the Q.a division in a *ham-1* mutant, a phenotype normally exhibited by Q.p. We found that this spindle movement in wild-type Q.p divisions required Ga proteins that promote spindle movement in other asymmetric divisions, and GPR-1, a protein involved in linking G proteins to microtubule asters, localized to the posterior cortex of Q.p. Genetic interactions suggest that *ham-1* mutant Q.a divisions also require Ga proteins function to divide with a reversed polarity. The transformation of Q.a to Q.p-like polarity in the *ham-1* mutant, however, appeared incomplete: *ham-1* loss did not alter the asymmetric localization of the non-muscle myosin NMY-2 to the anterior cortex of Q.a. A GFP tagged *ham-1* transgene revealed that Q.a but not Q.p expressed *ham-1*. Finally, we show that HAM-1 has both cortical and nuclear functions in Q,a DCSA. We propose a model where HAM-1 modifies a default Q.p-type polarity by localizing WASp function to the posterior Q.a membrane and by interfering with G-protein mediated spindle movement.

**Author Summary:** One way that animals produce different cell types is by asymmetric cell division, where a cell divides to produce daughter cells that differ in fate. Much is known about the mechanisms that polarize dividing cells to generate daughters that differ in fate. Some asymmetric cell divisions also result in daughters that differ in size, and the mechanisms that regulate how cells generate an asymmetric cleavage plane are poorly understood. In neural progenitors of the nematode *Caenorhabditis elegans*, two distinct mechanisms generate daughter cells of different size. One type requires movement of the mitotic spindle, which then defines the plane of the cell division. The other is spindle-independent. Here, we study two cells that divide with opposite polarities using these two mechanisms. We find that in the absence of the protein HAM-1, which has been reported to regulate gene transcription, the cell that normally divides using a spindle-independent mechanism now divides with a reversed polarity using a spindle-dependent mechanism. Our findings suggest that HAM-1 plays a key role in defining the mechanism by which a progenitor divides to produce daughter cells of different sizes and that both localization to the cell periphery and nucleus are important for its function.

## Introduction

The asymmetric division of progenitors to generate daughter cells of different fates leads to cellular diversity during metazoan development. Certain asymmetric cell divisions (ACDs) produce daughter cells that differ not only in fate but also in size. Many of the molecules that regulate ACD control both the asymmetric distribution of fate and the orientation of the mitotic spindle (for reviews of two models see [1, 2]). Some of these molecules also regulate divisions that produce daughter cells of different size, which we refer to here as Daughter Cell Size Asymmetry (DCSA). In screens for mutants with defects in apoptosis, investigators identified several molecules that control DCSA in the divisions that produce a smaller daughter cell that dies (reviewed in [3]). Correlation between loss of DCSA and loss of the apoptotic fate suggests that DCSA plays a role in apoptosis. However, two of these molecules, PIG-1 and HAM-1, also regulate DCSA in divisions that do not produce a daughter cell that dies [4, 5], suggesting that the role of these molecules in promoting the apoptotic fate is an indirect consequence of altering the size of the daughter cell fated to die.

The divisions of the Q.a and Q.p neuroblasts provide a paradigm for studying DCSA. These sister cells divide asymmetrically with opposite polarity to produce a smaller daughter cell that dies. Q.a produces a smaller anterior daughter cell (Q.aa), and Q.p produces a smaller posterior daughter (Q.pp). These cells divide using distinct DCSA mechanisms: Q.a divides by a spindle-independent, myosin-dependent mechanism, and Q.p divides by a spindle-dependent, myosin-independent mechanism [6]. The Q.a and Q.p divisions both require a PAR-4/PIG-1 kinase pathway and molecules involved in membrane trafficking [6-10], indicating that despite their distinct modes of division, there are underlying mechanisms that both divisions share.

HAM-1, by contrast, regulates Q.a but not Q.p DCSA [11]. Screens for mutants with defects in the migrations of the HSN neurons identified the first mutant alleles of *ham-1*[12], and it was later shown that the migration defect likely results from an earlier defect in the ACD that produces the HSN precursor [13, 14]. Loss of HAM-1 impairs a number of divisions that produce a smaller anterior cell that dies, including the division of Q.a [5, 11, 13, 14]. The HAM-1 protein contains a winged-helix domain known as a Storkhead box and localizes to both the nucleus and cortex [11, 13-15], and is reported to bind to and to transcriptionally regulate the gene *pig-1* [11].

In this report, we investigate the role of HAM-1 in the Q.a division. We found that loss of HAM-1 results in a reversal of Q.a polarity, causing it to divide like Q.p. Two processes that contribute to Q.a DCSA require HAM-1: the orientation of membrane extension to the posterior in a Wiscott-Aldrich protein (WASp)-dependent process and the suppression of G-protein-dependent posterior displacement of the mitotic spindle. By contrast, the Q.a-specific asymmetric distribution of the nonmuscle myosin NMY-2 was unaltered in a *ham-1* mutant, indicating that NMY-2 asymmetry is not sufficient for Q.a polarity. We found that Q.a DCSA requires both cortical and nuclear HAM-1 and discuss the role of HAM-1 is DCSA.

## Results

### *ham-1* loss reverses the polarity of Q.a DCSA

Phenotypic studies have shown that the gene *ham-1* primarily regulates asymmetric divisions that produce smaller anterior and larger posterior daughter cells [5]. In order to investigate how *ham-1* controls DCSA, we focused on the larval Q neuroblast lineage. The *C. elegans* Q lineage is experimentally attractive because it is accessible to imaging and because all of the differentiated cells of the lineage can be labeled in a single animal [6, 16]. The left and right Q lineages each produce three neurons and two cells that die [17]. Both the anterior daughter (Q.a) and posterior daughter (Q.p) cells of Q divide to produce an apoptotic daughter (Fig 1A). The Q.a division also produces the oxygen-sensing neuron A/PQR, and the Q.p division produces the precursor that divides to generate the mechanosensory neuron A/PVM and the interneuron SDQR/L. While both Q.a and Q.p produce daughter cells of different sizes with smaller daughter cells that die, the polarity of these neuroblasts are mirror images: the anterior daughter of Q.a (Q.aa) and the posterior daughter of Q.p (Q.pp) are the smaller daughter cells that die. The mechanisms that position the cleavage furrow asymmetrically differ in the Q.a and Q.p divisions: an asymmetric spindle displacement determines the position of the Q.p furrow, whereas the asymmetric distribution of the non-muscle myosin II NMY-2 determines the position of the Q.a furrow [6]. Like NMY-2, HAM-1 regulates DCSA and the apoptotic fate in the Q.a but not the Q.p, division [11].

**Fig 1.**
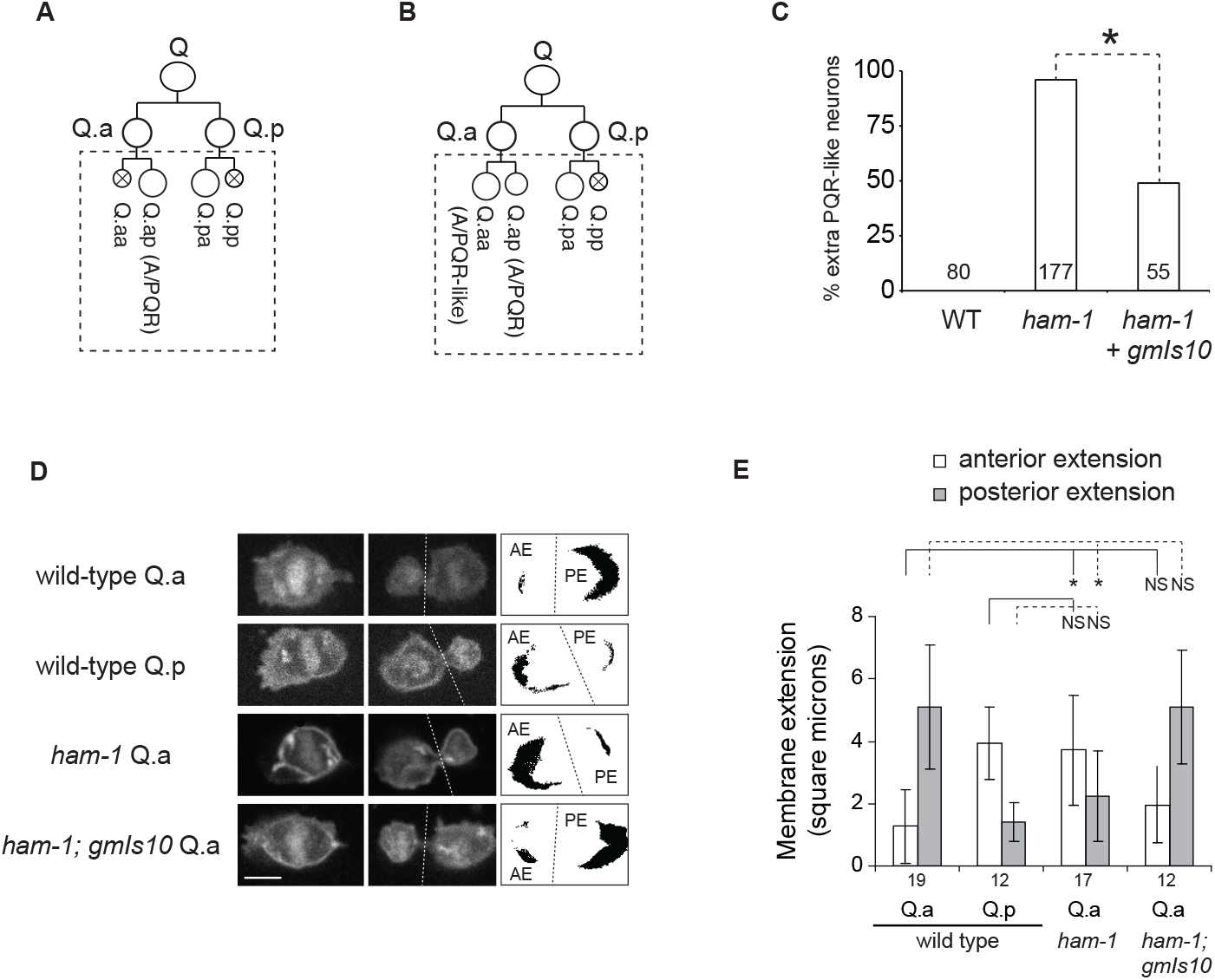
*ham-1* loss reverses the polarity of membrane extension in the Q.a neuroblast. Schematic diagrams of the Q lineage in wild-type (A) and *ham-1* (B) L1 larvae. *ham-1* mutations specifically impair Q.a DCSA leading to Q.aa survival and the production of extra A/PQR-like neurons. (C) Penetrance of the extra PQR-like neurons in wild-type and *ham-1* L1 larvae. The *gfp-ham-1* minigene *gmIs10* partially rescued the extra PQR-like neuron phenotype of *ham-1* mutants. The number of lineages analyzed is indicated over each bar. (D-E) Membrane extension measurements at the anterior and the posterior of dividing cells expressing the *rdvIs1[egl-17p::myr-mcherry; egl-17p::H2B::mcherry]*transgene, which labels the plasma membrane and chromosomes of the Q lineage cells. Anterior (AE) and posterior (PE) areas of extension are determined from metaphase (left panels) and telophase (center panels) time-lapse frames of Q.a and Q.p divisions after binarization and subtraction of the metaphase image from the telophase image (right panels). The dotted lines separate the anterior and posterior domains. Scale bar represents 3 µm. (E) Quantification of anterior and posterior membrane extension. The number of divisions analyzed is indicated under each bar * P<0.001 using the Student T-test; NS: not significant.

Using the Q lineage marker that labels each of the Q-derived neurons with a different color, we confirmed that *ham-1* mutations specifically perturb the Q.a division and lead to the production of extra A/PQR-like neurons (Fig S1). Q.a daughters of animals that bear mutations thought to either eliminate or strongly reduce *ham-1* function have a reversed DCSA with a larger Q.aa and smaller Q.pp (Fig 1B). Previous work led to the hypothesis that *ham-1* could act, at least in part, by controlling the transcription of the gene *pig-1* [11]. Another explanation is HAM-1 reverses a default Q.p polarity mechanism in Q.a. We tested these models.

### *pig-1* does not mediate the regulation of Q.a DCSA by *ham-1*

HAM-1 has been proposed to regulate the Q.a division by transcriptionally activating the gene *pig-1* [11]. The Q.a phenotypes of the *ham-1* and *pig-1* mutants are distinct: *ham-1* loss results in a reversed Q.a division, and *pig-1* loss results in a symmetric division. These phenotypic differences suggest that if HAM-1 transcriptionally activates *pig-1* to regulate Q.a DCSA, it must also regulate other genes. This proposed relationship between the two genes predicts that the *pig-1 ham-1* double mutant should have the same Q.a phenotype as a *ham-1* mutant. The *pig-1* single and *pig-1 ham-1* double mutants, however, produce similar Q.a DCSA phenotypes (Fig S2). The same epistatic interactions between *ham-1* and *pig-1* were previously observed for other lineages [8, 18].

The genetic interactions between *ham-1* and *pig-1* suggest that the *pig-1* gene may not be a functional HAM-1 target. To address the Feng et al. model directly, we estimated *pig-1* transcription levels in Q.a prior to its division in wild-type and *ham-1* mutants using a *pig-1p::gfp* transcriptional reporter (Fig S3). Although the GFP reporter showed a decreased expression in a strong SP1/*sptf-3* mutant as previously reported [19], none of the *ham-1* mutants tested showed a strong reduction in GFP expression. We measured a reduced GFP fluorescence intensity in the strong *ham-1(gm279)* and *ham-1(gm267)*mutants, but the weak *ham-1(cas46)* mutant previously studied by Feng et al. and the strong *ham-1(n1811)* mutant were not significantly different from wild-type animals.

One result consistent with HAM-1 acting as a regulator of *pig-1* is the ability of *ham-1* expression from the *egl-17* promoter to partially bypass the function of *ham-1* in the Q.a division [11]. To test if PIG-1 depletion at the post-transcriptional level could explain *ham-1* defects, we expressed a functional PIG-1::GFP translational fusion from the *mab-5* promoter in the left Q lineage and examined its ability to suppress the extra PQR-like cells defects in *ham-1* mutants. The complete rescue of the extra cell defects of *pig-1* null mutants confirmed that this fusion was functional in Q.a (Fig S4). We also confirmed that this fusion can partially suppress the defects of the *ham-1(cas46)* mutant, similar to the observation of Feng et al. using the *lin-17* promoter to express *pig-1*(Feng et al 2013). However, PIG-1::GFP expression did not suppress any of the other stronger mutants that we tested, suggesting that this effect is specific to the *cas46* allele or a difference in this mutant’s genetic background (Fig S4). Taken together, our findings suggest that the reduction of *pig-1* expression does not explain the ACD defects of *ham-1* mutants.

### *ham-1* loss reverses the orientation of Q.a membrane extension

To further investigate the possible functions of *ham-1* in Q.a DCSA, we followed the Q.a and Q.p divisions of wild-type animals, *ham-1* mutants, and *ham-1* mutants rescued with a *ham-1* transgene, *gmIs10*, which is a *ham-1* transgene tagged near the N-terminus with GFP. This trasgene was able to rescue partially the production of extra PQR-like neurons of *ham-1* mutants (Fig 1C). The dividing Q.a and Q.p neuroblasts extended their plasma membranes in opposite directions during anaphase: Q.a extended its membrane toward the posterior, which Ou et al. previously reported [6], and we report here that Q.p also extended its membrane, but in the opposite direction toward the anterior (Fig 1D, E). Strong *ham-1* mutations reversed the direction of Q.a membrane extension from anterior to posterior, the direction of membrane extension seen in dividing wild-type Q.p neuroblasts. The *gmIs10* transgene rescued this defect (Fig 1D, E).

WSP-1 is the *C. elegans* homolog of the Wiskott-Aldrich Syndrome Protein, which functions to nucleate the Arp2/3 complex and is activated by the Rho-family GTPase Cdc42 (for a review see [20]. A GFP-tagged WSP-1 localizes near the leading edges of the Q.a and Q.p neuroblasts during their migrations [21]. We found that the endogenously GFP-tagged gene *wsp-1* used in the migration study also localized to the extending membranes of Q.a and Q.p during their divisions, and that the localization of WSP-1::GFP was reversed in *ham-1* mutant Q.a neuroblasts (Fig 2A-E).

**Fig 2.**
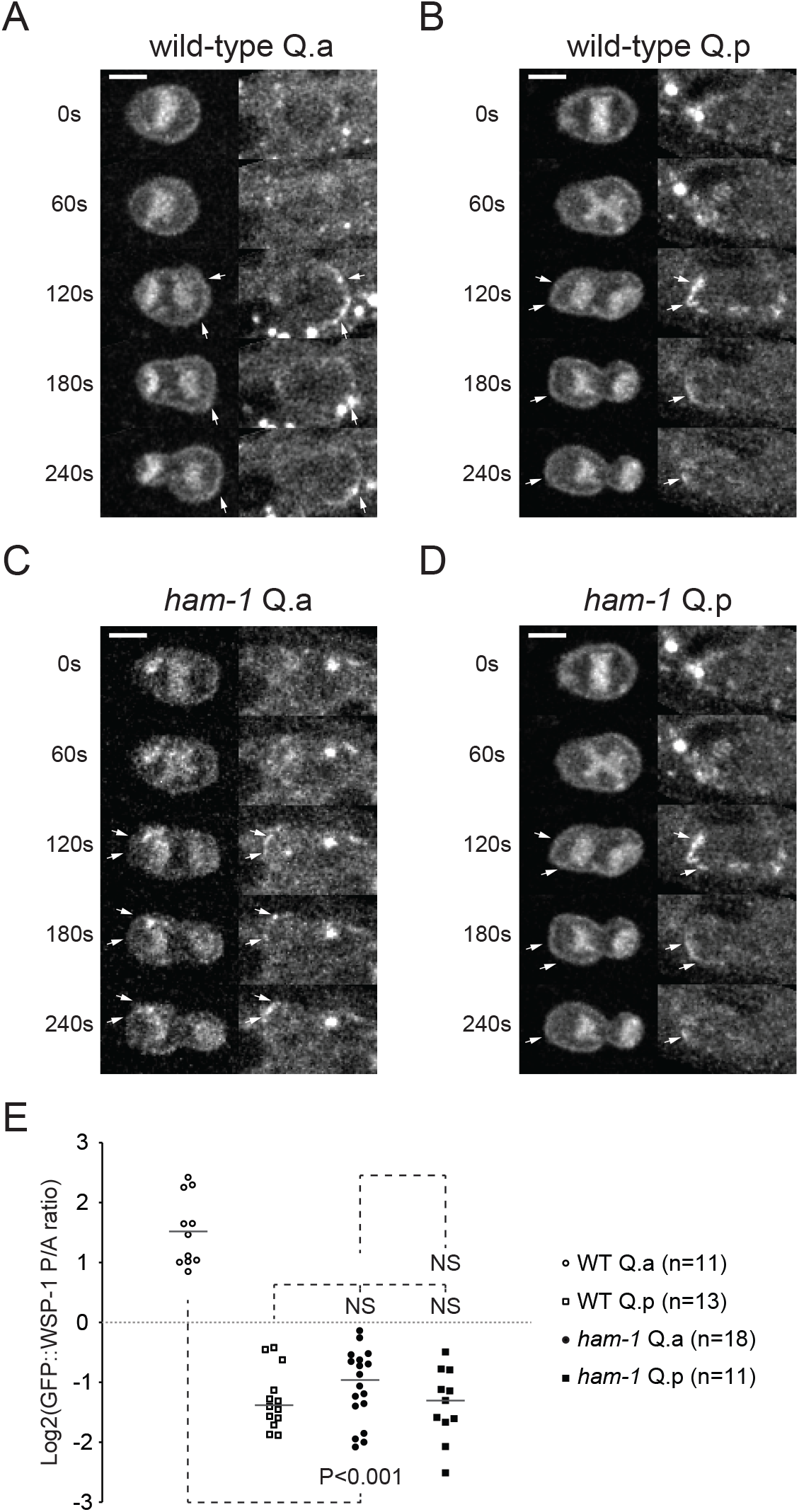
WSP-1 localizes at the extending plasma membrane of Q.a and Q.p in anaphase. (A-D) Confocal time lapse imaging of GFP::WSP-1 localization in dividing QR.a (A, C) and QR.p (B, D) cells in wild-type (A, B) and *ham-1* (C, D) L1 larvae carrying *casIs165[egl-17p::myr-mCherry; egl-17p::mCherry-TEV-S]*which labels the plasma membrane and the chromosomes of the Q cells (left panels), and *cas723[gfp::wsp-1a knock-in]*which labels the actin regulator WSP-1 (right panels) [21]. White arrows indicate membrane localization of WSP-1 (right panels) and the corresponding labeled membranes (left panels). Time is indicated in seconds post-chromosome separation (t=0s). Scale bars represent 3 µm. (E) Distributions of WSP-1::GFP intensity ratios of the posterior and the anterior membranes at anaphase (t=180s after chromosome separation) for the Q.a and Q.p cells of wild-type and *ham-1* L1 larvae. P-values obtained using the Mann-Whitney U-test are indicated; NS: not significant.

The WSP-1::GFP localization implies that the protein functions in the Q.a and Q.p divisions. To test this hypothesis, we measured Q.a and Q.p daughter cell sizes in a *wsp-1* mutant, as well as in the *cdc-42* and Arp2/3 mutants *arx-2* and*arx-7*. Although all of the mutants displayed reduced Q.a and Q.p DCSA, only the *wsp-1* and *arx-2* mutants exhibited a significant reduction in Q.p DCSA (Fig S5). These findings support the model that the *ham-1* mutant Q.a adopts a Q.p mechanism of membrane extension dependent on WASp and the Arp2/3 complex.

### *ham-1* loss results in a G-protein mediated Q.a spindle displacement

We also tracked spindle movement by kymographic analysis of the Q.a and Q.p anaphase chromosomes. As described previously [6], the posterior chromosomes moved further than the anterior chromosomes during wild-type Q.p divisions, and both sets of chromosomes moved symmetrically toward the poles during wild-type Q.a divisions. As in the wild-type Q.p division, chromosome displacement during the Q.a division of *ham*-*1* mutants was similarly skewed towards the posterior of the cell, and this defect was rescued by the *gmIs10* transgene (Fig 3A,B).

**Fig 3.**
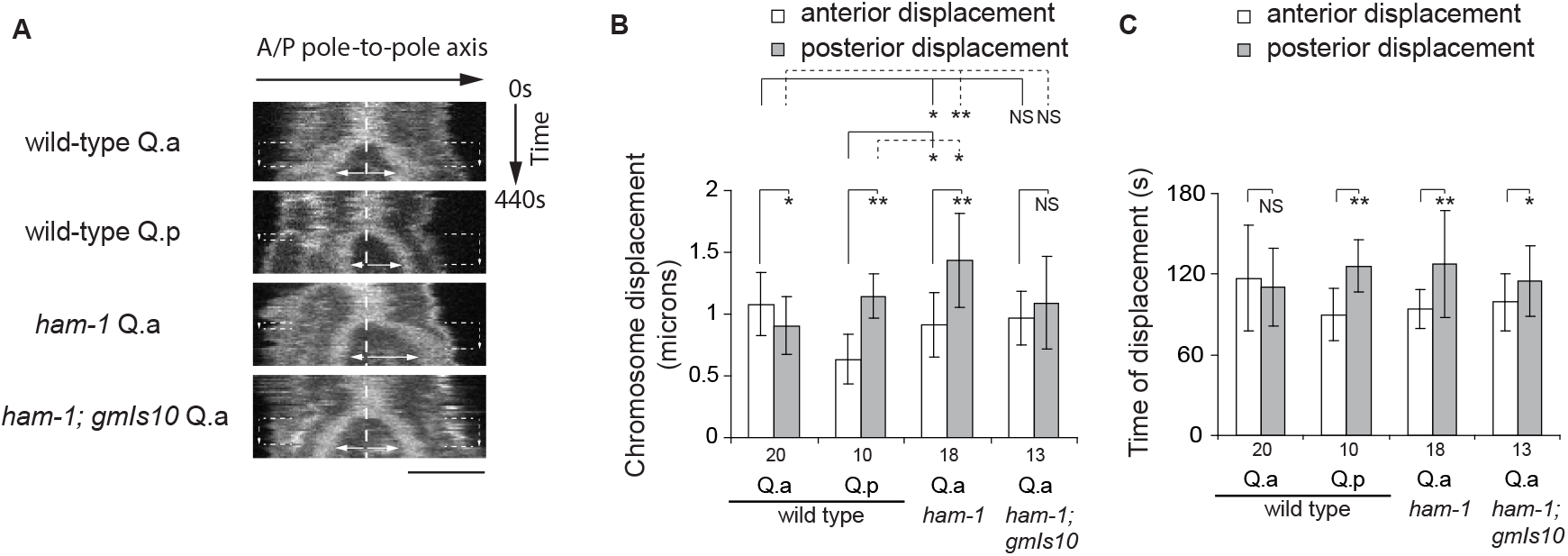
The Q.a mitotic spindle in *ham-1* mutants undergoes a Q.p-like posterior displacement. (A) Kymographs of chromosome displacement during anaphase from time-lapse imaging of Q.a and Q.p divisions in wild-type animals, and Q.a divisions in *ham-1* and rescued *ham-1; gmIs10* mutant L1 larvae. Each kymograph was compiled from 44 images taken at 10 s intervals (y axis). The vertical dashed line indicates the center of the metaphase plate in the last frame of metaphase. Horizontal arrows indicate the length of maximum chromosome displacement towards the anterior and the posterior poles. The vertical arrows between horizontal dashed lines indicate the duration of anterior and posterior displacements. Scale bar: 3 µm. (B) Average anterior and posterior chromosome displacement. (C) Average duration of anterior and posterior chromosome displacement. In (B) and (C), numbers under the bars are the number of divisions scored and error bars represent standard deviations. ** P<0.001, * P<0.05, using the Student T-test; NS not significant.

Our kymographic analysis of chromosome movement also revealed that the timing of chromosome movement differed in the Q.a and Q.p divisions. The anterior and posterior sets of wild-type Q.a chromosomes were displaced toward the cell poles for similar amounts of time. The movement of the posterior chromosomes from wild-type Q.p and *ham-1* mutant Q.a, by contrast, continued after movement of the anterior chromosomes, timing defects that were partially rescued by the *gmIs10* transgene (Fig 3C). These measurements suggest that Q.p regulates furrow positioning by arresting the anterior chromosome movement and by prolonging posterior chromosome movement.

Because G proteins regulate spindle orientation and positioning in the *C. elegans* zygote and Drosophila neuroblasts (for reviews see [1, 2]), they are attractive candidates to be regulators of spindle-dependent Q.p DCSA. In the *C. elegans* zygote, two Gα proteins, GPA-16 and GOA-1, function in parallel as monomeric GDP-bound subunits to recruit the GoLoco proteins GPR-1/2 and microtubule-dependent force generators to the cortex, leading to mitotic spindle displacement (reviewed in [2]). Because the *gpa-1* and *goa-1* mutations result in a maternal-effect lethal phenotype, we were unable to analyze mutants that lacked both zygotic and maternal function. For *gpa-16*, we shifted the temperature-sensitive *gpa-16(it143)* mutant to the restrictive temperature at hatching and observed a weak but significant DCSA reduction only for Q.p (Fig 4A). By analogy to the *C. elegans* zygote, the lack of a strong Q.p DCSA defect in *gpa-16(it143)* mutant larvae could result from functional compensation by other Gα proteins. For *goa-1*, we analyzed animals that lacked zygotic but not maternal function and found weak effects on DCSA in both Q.a and Q.p (Fig 4A,B). Because of lethality, we were unable to analyze the double mutant or animals where one gene was mutated and the function of the other reduced by RNAi.

**Fig 4.**
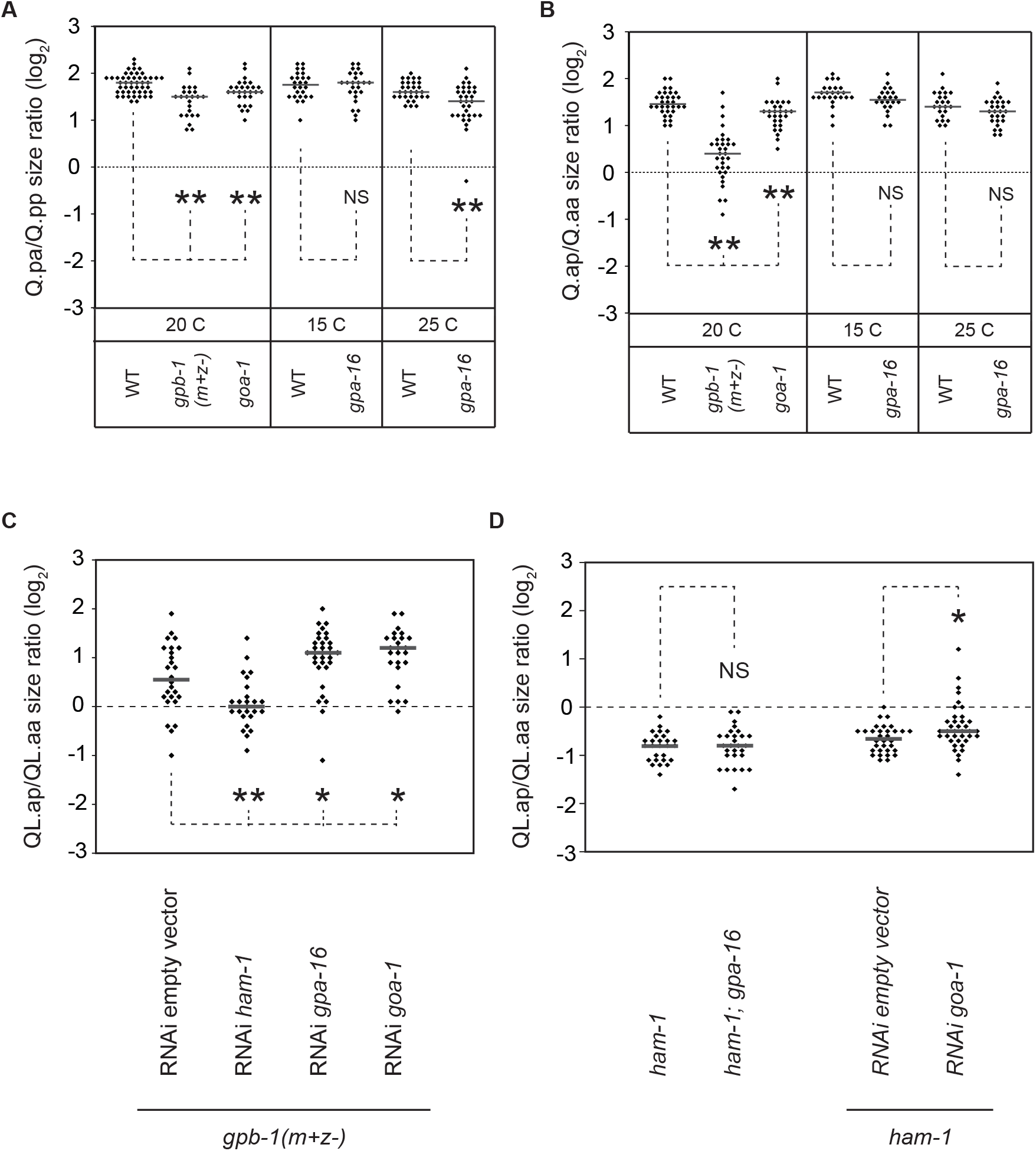
Trimeric G proteins regulate Q.p and Q.a DCSA. (A-D) Distribution of cell-size ratios of Q.p (A) and Q.a (B-D) daughters of wild-type and trimeric G proteins mutant L1 larvae. Cells were labelled using the *gmIs81[egl-17p::gfp]* marker. (A, B) The *gpa-16(it143)* thermosensitive mutant and the control strain were scored at permissive (15°C) and restrictive (25°C) temperatures. (C) Distribution of cell size ratios between Q.a daughters in *gpb-1(m+z-)* L1 larvae fed with bacterial clones carrying *C. elegans* genomic DNA sequences cloned into the L4440 plasmid. These bacteria generate double-stranded RNA upon IPTG induction. The L4440 empty vector was used as negative control. A L4440-*ham-1* clone was used as positive control. (D) Distribution of cell-size ratios between Q.a daughters in *ham-1* single mutant and *ham-1; gpa-16* double mutant L1 larvae grown at semi-permissive temperature (20°C), and in *ham-1* single mutant L1 larvae fed with bacterial RNAi clones. For each RNAi condition, data points were cumulated from at least three independent experiments. ** P<0.01, * P<0.05 using the Mann-Whitney U-test; NS: not significant.

Mutant animals lacking zygotic but not maternal function of the Gβ gene *gpb-1* can escape embryonic lethality and arrest during the L1 stage (Zwaal et al 1996), allowing us to examine the Q lineage. Q.p DCSA of *gpb-1* larvae was only slightly decreased, whereas the Q.a DCSA was more severely affected (Fig 4A,B). One possible explanation for the later observation is that Gβγ acts directly to promote Q.a DCSA, independent of Gα. Alternatively, the Q.a phenotype could result indirectly from the increased Gα-GDP levels. In the *C. elegans* zygote, the formation of a trimeric complex inhibits Gα-GDP activity, and reduction of the Gβ GPB-1 function increases monomeric Gα-GDP subunits, which promote spindle movement [22]. This later explanation for the *gpb-1* effects predicts that aberrant Gα activity in a *gpb-1* mutant shifts the Q.a spindle to the posterior. Because other mechanisms that promote Q.a asymmetry would be intact, for example, posterior membrane extension, the posterior shift of the spindle would lead to a more symmetric division. This model predicts that reduction of Gα function in a *gpb-1* mutant will lead to a more asymmetric division that is more like a wild-type Q.a DCSA. Consistent with this hypothesis, reduction of *goa-1* or *gpa-16* by RNAi in a *gpb-1* mutant background resulted in a more asymmetric Q.a division (Fig 4C), supporting the model that a *gpb-1* induced phenotype results from a gain of Gα function. *ham-1* RNAi was used as a positive control for RNAi and significantly enhanced the *gpb-1* mutant DCSA phenotype (Fig 4C).

If the *ham-1* Q.a neuroblasts adopt a Q.p division pattern, then reducing Gα signaling in a *ham-1* mutant should disrupt DCSA. To test this prediction, we used the *goa-1* and *gpa-16* mutations to reduce their function in the *ham-1(gm279)* mutant and observed a weak but significant reduction in DCSA with the *goa-1* but not the *gpa-16* mutation (Fig 4D).

In the zygote, the two closely related TPR-GoLoco paralogs, GPR-1 and GPR-2, are required for spindle asymmetry. Because TPR-GoLoco proteins interact with GDP-bound Gα proteins, they can be used as biosensors for the distribution of GDP-Gαs [22-25]. As in the early embryo [26], mCherry::GPR-1 localized to the centrosomes of Q.a and Q.p during anaphase (Fig 5A,B). At the start of anaphase, mCherry::GPR-1 accumulated at the posterior cortex of both wild-type and *ham-1* mutant Q.p cells, but did not localize asymmetrically during the Q.a division (Fig 5A,B,D,E). The posterior movement of the chromosomes in the mutant *ham-1* Q.a division suggests that mCherry::GPR-1 would localize at the posterior in mutant *ham-1* Q.a during anaphase divisions. Although there was a difference in the distribution of the mCherry::GPR-1 in mutant *ham-1* Q.a cells, the shift in distribution was not significant, perhaps due to limitations of the reporter to detect low levels of Gα activation (Fig 5C-E). Taken together, our findings suggest that HAM-1 has two functions in Q.a, repressing G-protein mediated posterior spindle movement and maintaining WSP-1-dependent membrane extension at the posterior.

**Fig 5.**
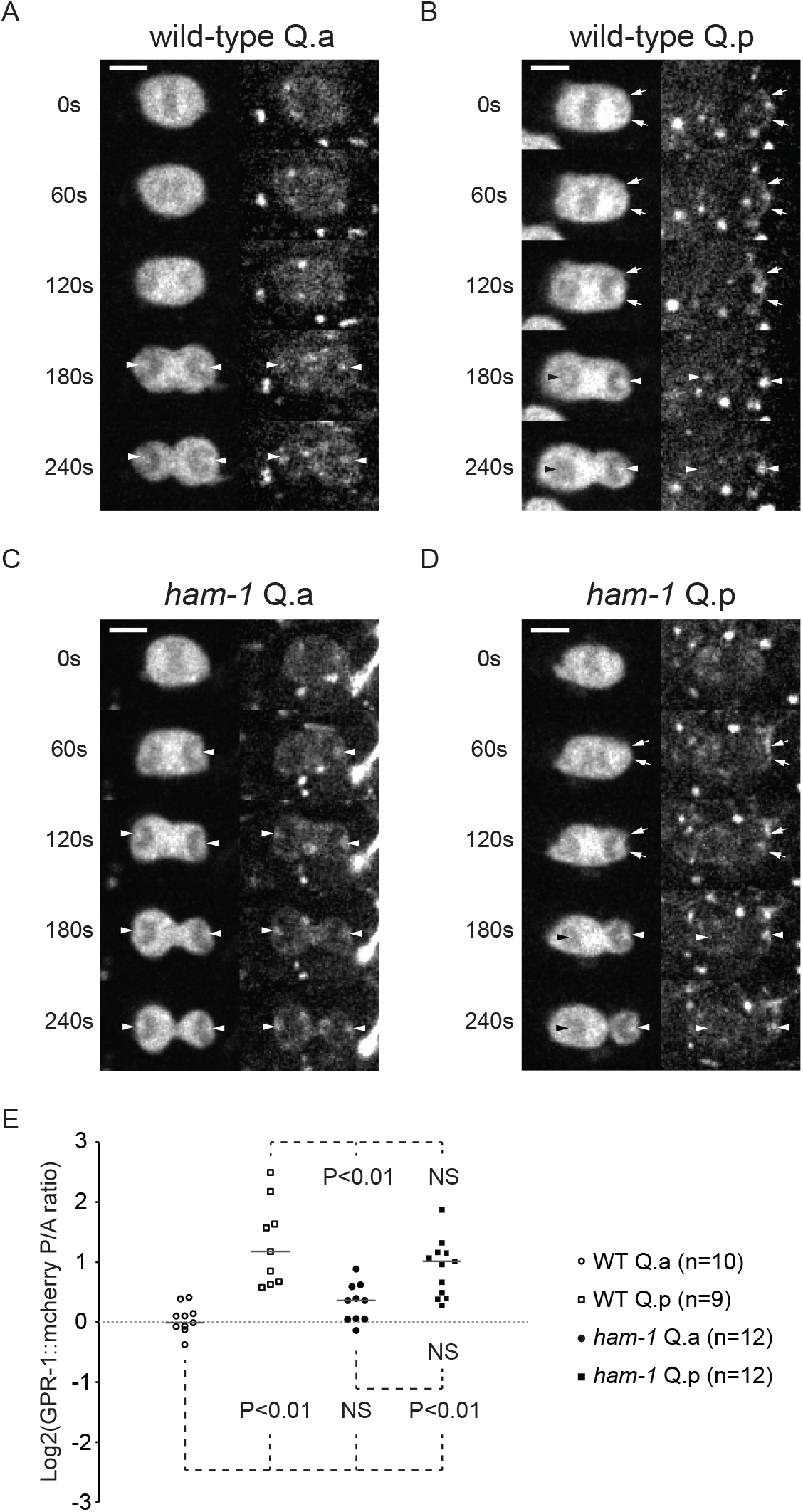
GPR-1 transiently localizes at the posterior pole of dividing Q.p at the start of anaphase. (A-D) Confocal time-lapse imaging of GPR-1::mcherry localization in dividing QR.a (A, C) and QR.p (B, D) cells of wild-type (A, B) and *ham-1* (C, D) L1 larvae carrying the *gmIs81[egl-17p::gfp]*transgene, which labels the Q lineage cells, and the *ddIs79[mCherry::gpr-1]* transgene, which labels the GoLOCO protein GPR-1 which binds to free GDP-bound Gai/o (right panels) [26]. Arrows indicate membrane localization of GPR-1 (right panels) and the corresponding labeled membranes (left panels). Arrowheads indicate GPR-1 localization at the centrosome. Time is indicated in seconds post-chromosome separation (t=0s). Scale bars represent 3 µm. (E) Distributions of GPR-1::mcherry intensity ratios between the posterior and the anterior membranes in early anaphase (t=20s after chromosome separation) for Q.a and Q.p cells of wild-type and *ham-1* L1 larvae. P-values obtained using the Mann-Whitney U-test are indicated; NS: not significant.

### Loss of *ham-1* function does not alter NMY-2::GFP distribution in Q.a

A GFP-tagged nonmuscle myosin II NMY-2 is asymmetrically distributed to the anterior of Q.a, the pole that does not extend during anaphase, but is symmetrically distributed in Q.p [6]. Ou et al. also showed that Chromaphore Assisted Laser Inactivation (CALI) of the anteriorly distributed NMY-2::GFP disrupted Q.a DCSA, supporting the hypothesis that asymmetric NMY-2 activity restricts membrane extension to the posterior of Q.a [6]. This mechanism appears to be conserved in Drosophila neuroblasts, where the myosin light chain Spaghetti Squash (Sqh) transiently localizes asymmetrically to the side of the neuroblast that will produce the smaller daughter, the ganglion mother cell [27, 28]. Sqh asymmetry occurs at the pole opposite of the neuroblast extension, and mutations that disrupt the localization of Sqh result in cortical extension at both poles [27, 28].

The model that Q.a adopts the Q.p mechanism of asymmetric division when *ham-1* function is lost predicts that *ham-1* loss should also result in a loss of NMY-2 asymmetry in Q.a. Feng et al. confirmed this prediction using a weak *ham-1* allele [11]. We repeated this experiment using the same *nmy-2::gfp* transgene and *gm279*, a nonsense allele of *ham-1* that results in no detectable HAM-1 protein [13, 14], and were surprised to find that NMY-2::GFP still localized asymmetrically to the anterior of Q.a (Fig 6A-C). It is clear that this localization occurs in a *ham-1* mutant background because following the division shows that DCSA is reversed and NMY-2::GFP localized to the side of the Q.a that will produce the larger daughter cell. This observation leads to the surprising conclusion that the asymmetric distribution of NMY-2 is not sufficient for Q.a polarity.

**Fig 6.**
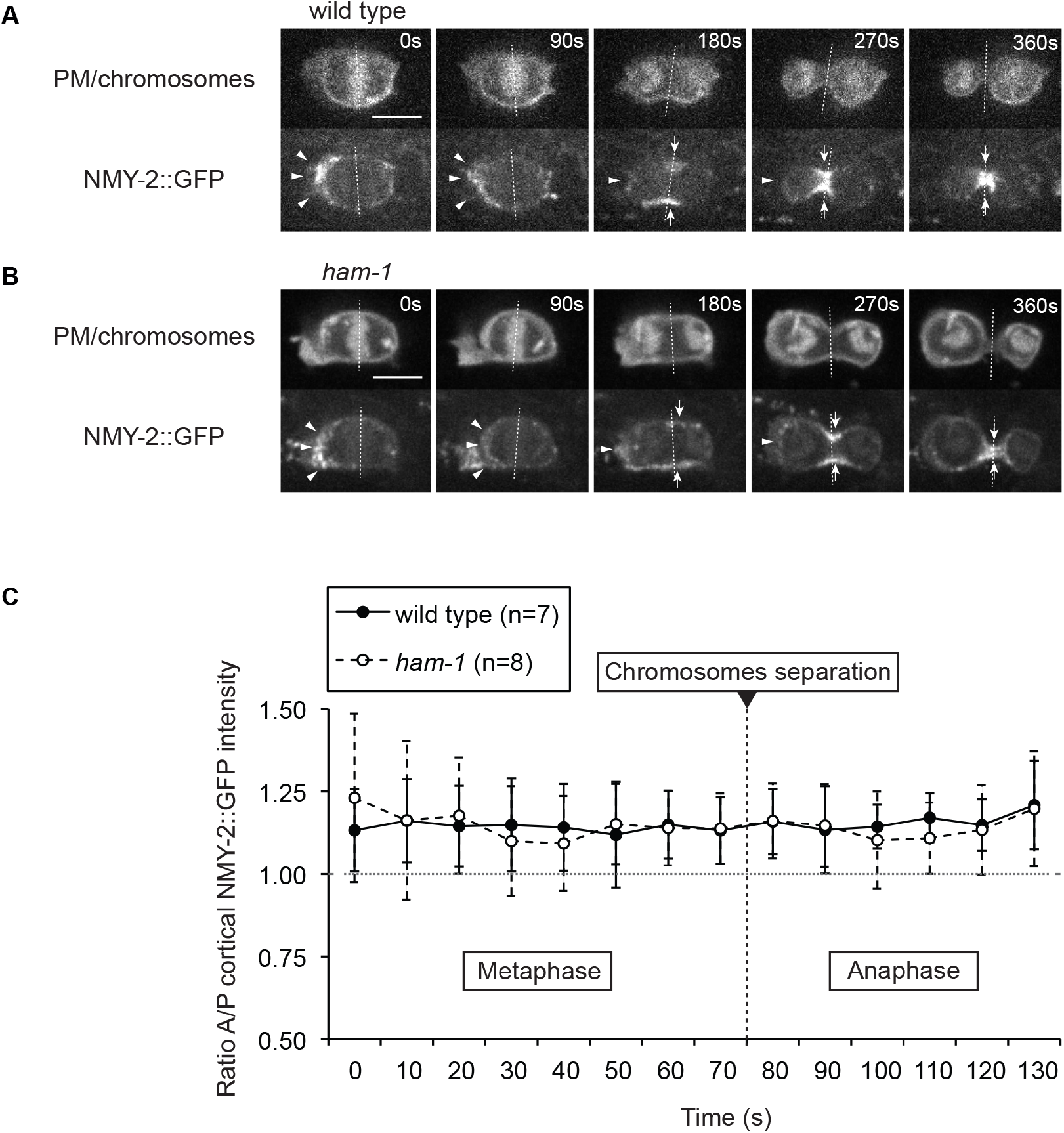
Loss of *ham-1* does not alter NMY-2::GFP polarity. (A, B) Time-lapse imaging of NMY-2::GFP localization in the dividing QR.a cell of (A) wild-type and (B) *ham-1(gm279)* L1 larvae carrying *rdvIs1[egl-17p::myristoylated mcherry; egl-17p::H2B::mcherry]*, which labels the plasma membrane and chromosomes of the Q lineage cells, and the *zuIs45[nmy-2p::nmy-2::gfp]* transgene, which labels the myosin heavy chain NMY-2. White arrowheads indicate anterior accumulation of cortical myosin during metaphase and early anaphase. White arrows indicate myosin at the cleavage furrow during cytokinesis. Scale bars represent 3 µm. Dotted lines demark the anterior posterior border as defined in the Materials and Methods section. (C) The anterior/posterior (A/P) NMY-2::GFP intensity ratios from the last eight frames of metaphase and first six frames of anaphase (before the formation of the cleavage furrow) in wild-type and *ham-1(gm279)* mutant L1 larvae. The transition from metaphase to anaphase is defined by chromosomes separation. The horizontal dotted line shows 1:1 AP intensity ratios. Error bars represent standard deviations. At each time point, NMY-2::GFP intensity ratios show that there was no significant difference in the anterior myosin polarity between wild type and *ham-1* cells.

### Q.a but not Q.p expresses *ham-1*

Transgenic expression of a translational GFP::HAM-1 fusion under the control of the endogenous *ham-1* 5', intronic and 3' regulatory sequences (*gmIs10*) rescued a subset of *ham-1* lineage defects (data not shown), and also partially rescued the extra A/PQR-like phenotype of *ham-1* mutants (Fig 1C). Surprisingly, the ModENCODE transgene *wgIs102 [Pham-1::ham-1::gfp]*shown by Feng et al. to partially rescue the hypomorphic allele *ham-1(cas46)* failed in our hands to rescue the extra neuron defects of strong *ham-1* mutants (data not shown).

HAM-1 protein was initially detected at the cell cortex of dividing embryonic cells using anti-HAM-1 antibodies [14]. Using GFP-tagged HAM-1, two groups detected nuclear HAM-1 [11, 15]. The presence of endogenous HAM-1 in the nucleus was confirmed by cell fractionation and immunoblotting with anti-HAM-1 antibodies that detect only cortical HAM-1 in stained embryos [15]. Leung et al. used a GFP::HAM-1 fusion expressed from a heterologous promoter, precluding analysis of the *ham-1* expression pattern [15]. The *wgIs102* transgene used by Feng et al., by contrast, should contain the full-length *ham-1* gene with GFP fused to the C-terminus of the protein [11, 15]. The *wgIs102* transgene revealed broad expression of GFP in somatic nuclei, including all cells of the Q lineage, and did not localize to the cortex [11]. We also observed this GFP expression and localization pattern with the *wgIs102* transgene, which differed from the more dynamic expression pattern and cortical localization of GFP::HAM-1 from animals containing the *gmIs10* transgene. For example, we detected GFP::HAM-1 in Q.a and its daughter cells, but not in the Q.p lineage. GFP::HAM-1 localization is also dynamic in the Q.a lineage: in Q.a, it localizes to the cortex during mitosis, but to the nucleus before nuclear envelope breakdown. After cytokinesis, GFP::HAM-1 is visible in both Q.a daughter nuclei (Fig 7A).

**Fig 7.**
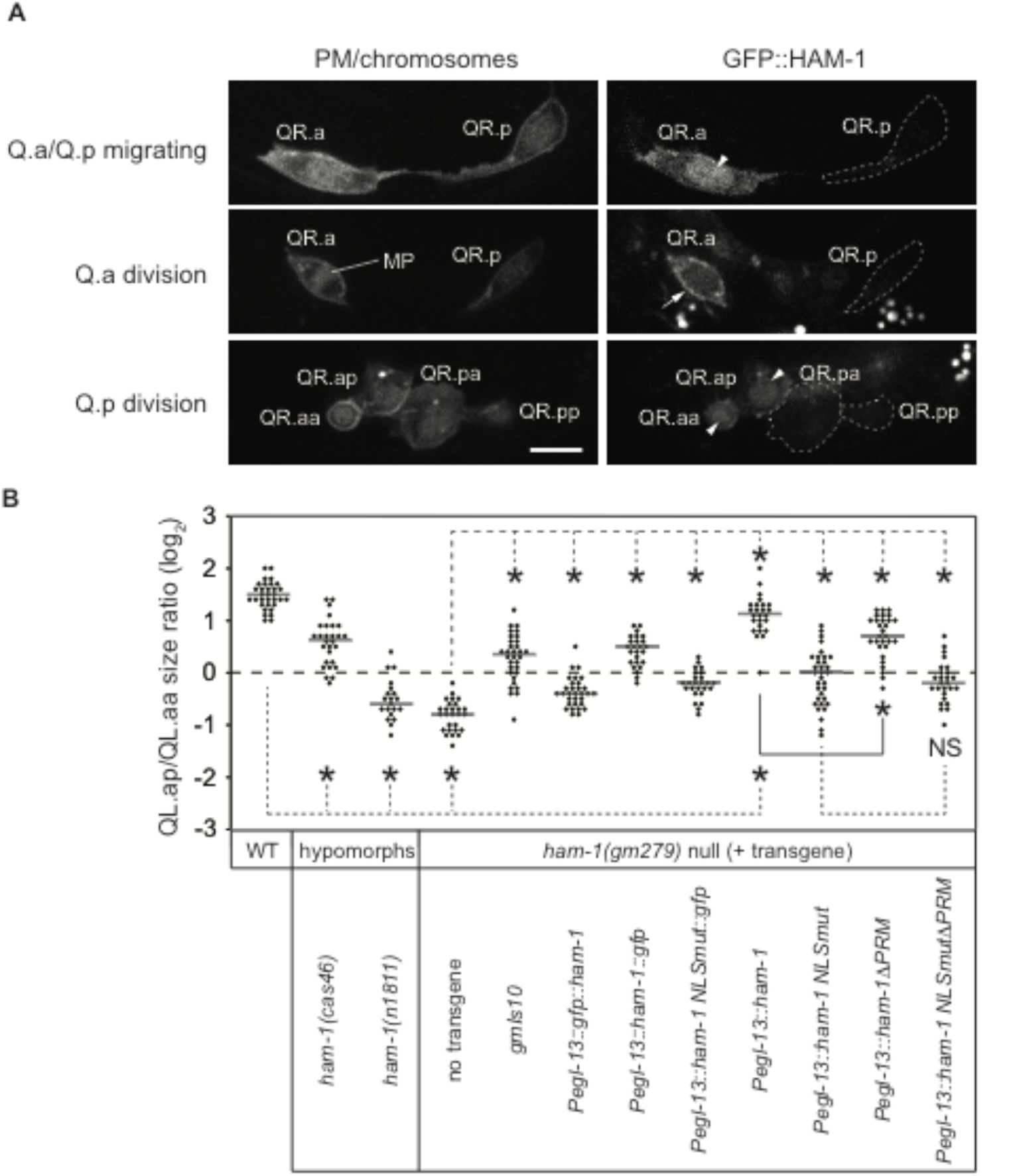
HAM-1 functions autonomously in Q.a in the nucleus and at the plasma membrane. (A) GFP::HAM-1 expression and localization in the Q lineage. Representative confocal micrographs showing plasma membrane (PM) and chromosomes (left panels) and GFP::HAM-1 expression (right panels) for each stage of the QR lineage after Q division in *ham-1(gm279)* animals carrying the *rdvIs1* and the rescuing *gmIs10[gfp::ham-1]* markers. For all photomicrographs, anterior is to the left. (Q.a/Q.p migration) The right Q.a and Q.p migrate anteriorly. On the left side (not shown), Q.p does not migrate, but Q.a migrates posteriorly. (Q.a division and Q.p division) A chromosomal metaphase plate (MP) is visible in Q.a which divides before Q.p. White arrows and arrowheads indicate a Q.a-specific cortical and nuclear localization of GFP::HAM-1, respectively. Similar GFP::HAM-1 localizations were observed in the QL.a lineage (data not shown). Scale bar represents 3 µm. (B) Cell size ratios between QL.a daughters, labelled by the *gmIs81[egl-17p::gfp]*marker, in wild-type, *ham-1* and *ham-1* transgenic L1 larvae. The DCSA is lost and progressively reversed in an allelic series of *ham-1* mutants. The *gmIs10[gfp::ham-1]*transgene partially restores DCSA in the *ham-1(gm279)* null background. Using single-copy MosSCI transgenes, GFP-tagged and untagged forms of *ham-1* were expressed in Q.a, partially rescuing DCSA as well. * P<0.001 using the Mann-Whitney U-test, NS: not significant.

The Q.a-specific expression of our *gmIs10* transgene is consistent with the requirement for *ham-1* function in Q.a but not Q.p. The cortical localization of GFP::HAM-1 in these transgenic animals is also consistent with previous anti-HAM-1 antibody staining and the cortical localization of HAM-1::GFP [13-15]. To determine the basis for the discrepancy between the *gmIs10* and *wgIs102* transgenes, we carried out an inverse PCR analysis of genomic DNA from a strain containing *wgIs102* and found that *wgIs102* is a *mep-1::gfp* transgene, incorrectly annotated as a *ham-1::gfp* reporter (data not shown). The expression of GFP from *wgIs102* is consistent with the expression pattern and localization of the zinc-finger protein MEP-1 [29].

The previous description of the *ham-1* expression pattern was carried out in embryos by antibody staining [13, 14]. We used *gmIs10* to further characterize *ham-1* postembryonic expression pattern. We observed nuclear and cortical GFP::HAM-1 expression in the hypodermal seam cells, in cells of the V5.pa lineage and in unidentified neurons (data not shown). We detected GFP::HAM-1 in daughters of V5.paa, consistent with its function in regulating DCSA for this division [5]. By contrast, a survey of the seam cell divisions in a *ham-1* mutant revealed no obvious role for HAM-1 (data not shown).

### *ham-1* acts autonomously in the Q.a neuroblast

The *gmIs10* reporter indicates that HAM-1 is expressed in Q.a and in cells that are nearby, for example, the seam cells. To test whether HAM-1 functions in Q.a to regulate its ACD, we created transgenes that drive HAM-1 expression from the Q.a-specific *egl-13* promoter [11]. To be able to compare different transgenes and to reduce the effects of excess expression, we constructed single-copy transgenes integrated at the same genomic locus using the Mos1-Mediated Single-Copy Insertion (MosSCI) technique [30]. These MosSCI transgenes were then introduced into *ham-1* mutants. Their ability to rescue the *ham-1* cell size reversal and extra neuron defects was assessed using the *gmIs81* reporter that labels the dividing neuroblasts and their final neuronal descendants using distinct fluorescent proteins [16].

Unexpectedly, MosSCI trangene-driven expression of an N-terminally or C-terminally GFP-tagged HAM-1 fusions in Q.a either failed to rescue or rescued poorly the cell size reversal defects of the *ham-1* mutants. However, an untagged HAM-1 protein rescued most Q.a divisions (Fig 7B). The extra A/PQR neuron defects showed a similar pattern of rescue with independently derived transgenes (Fig S6). These results extend the finding that HAM-1 functions autonomously in the Q lineage [11], showing that HAM-1 is expressed and acts in Q.a. Our data also indicate that tagging of HAM-1 at either end impairs its function. This perturbation of function by tagging may explain why membrane localization of the GFP::HAM-1 fusion during mitosis is continuous around the cell, and different from the posterior accumulation of endogenous HAM-1 previously revealed by antibody staining [13, 14].

### HAM-1 appears to act both in the nucleus and at the cortex

Two Nuclear Localization Sequences (NLSs) have been shown to control HAM-1 nuclear import [15]. In an attempt to determine whether HAM-1 acts in the nucleus, both NLSs were mutated in MosSCI-generated transgenes that express HAM-1::GFP and untagged HAM-1 proteins. Expression of the untagged, NLS-mutant proteins in Q.a partially rescued *ham-1* phenotypes (Fig 7B), suggesting that nuclear localization is partially required for full HAM-1 function. The GFP-tagged proteins failed to localize to the nucleus (data not shown).

Several *ham-1* alleles have been shown to disrupt or decrease HAM-1 cortical localization. For instance, the *ham-1(n1811)* mutation results in a G47D change in the N-terminus of HAM-1 and eliminates its cortical accumulation in embryos [14]. In the PLM/ALN neuroblast lineage, *ham-1(gm279)* mutants display a high penetrance of missing neurons, but *ham-1(n1811)* mutants display a mild phenotype, suggesting that HAM-1 cortical localization is not critical for the ALN/PLM lineage [15]. By contrast, *ham-1(n1811)*mutant animals display severe Q.a defects (Fig 7B), suggesting that Q.a DCSA requires both nuclear and cortical HAM-1 functions. The other mutations in the N-terminus of HAM-1 also reduce its accumulation at the cortex and have significant Q.a defects (Fig 7B)

Leung et al. also showed that mutating a proline-rich motif (PRM), a putative SH3-binding domain, drastically reduced GFP::HAM-1 cortical localization (Leung et al 2015). Expression of an untagged version of this mutant reduced the ability of the transgene to rescue the *ham-1* mutant Q.a DCSA defects when compared to the untagged wild-type transgene (Fig 7B). Combining the NLS and PRM mutations resulted in less rescue of the *ham-1* DCSA defect, although the difference between the NLS and the combined NLS and PRM mutant rescue was not significant (Fig 7B). However, the ability of these two transgenes to rescue the apoptosis defect of *ham-1(gm279)*mutants was significantly different: the NLS mutant had 69% extra PQR-like neurons, the PRM mutant had 6% extra PQR-like neurons, and the double NLS, PRM mutant had 90% extra PQR-like neurons, a phenotype similar to the non-transgenic *ham-1* mutant (Fig S6). This latter finding could mean that HAM-1 has an apoptotic function that depends on the PRM, an activity that is separable from its role in DCSA. Alternatively, rescue of the apoptosis defect is a more sensitive readout of subtle DCSA defects. Taken together, our findings suggest that HAM-1 requires both nuclear and cortical functions to control Q.a DCSA.

### Correlation between the *ham-1* DCSA and apoptosis defects

The analysis of various rescuing constructs and *ham-1* alleles discussed above provided us with a range of daughter-cell size ratios from normal to reversed. When plotted against the percentage of extra PQR neurons observed in the same strains, we observed a clear correlation between a reversal in size of QL.a daughter cell sizes and the ability of QL.aa to survive and adopt a PQR-like neuron fate (Fig S7). A similar correlation between increased cell size and survival was previously observed for Q.pp [3] and the NSM neuron's apoptotic sister [31]. A more recent study suggests that CED-3 caspase activity is asymmetrically distributed in divisions that produce an apoptotic daughter cell [32]. The correlation between the apoptotic fate and daughter cell size support the model where the reduction of cell size by ACD also promotes the apoptotic fate by increasing the concentration of CED-3 in the smaller daughter.

## Discussion

### HAM-1 promotes Q.a membrane extension but not myosin localization

Ou. et al originally showed that posterior membrane extension contributed to the asymmetry of the Q.a division [6]. We confirmed this finding and showed that anterior membrane extension also contributes to the asymmetry of the Q.p division. Localization of WSP-1::GFP and the effects of *wsp-1, cdc-42*, *arx-2* and *arx-7* mutations suggest that polarized actin polymerization promotes membrane extension for both divisions. The Q.a and Q.p cells also migrate, and both WSP-1 and the *C. elegans* WAVE homolog WVE-1 act in parallel to promote the extension of the leading edge of these migrating cells [21]. The specificity of the *ham-1* mutations for the Q.a is unusual as most of the mutations that alter DCSA in the Q lineage affect both the Q.a and Q.p divisions [6, 8-10]. Because membrane extension contributes to DCSA in both Q.a and Q.p, these genes may regulate the molecules that promote membrane extension.

Our findings and those of Feng et al. indicate that *ham-1* mutations alter the Q.a, but not the Q.p division [11], and our analysis of strong *ham-1* mutants showed that the Q.a divisions were reversed and behaved in several ways like wild-type Q.p divisions when *ham-1* function is eliminated. The reversal of WSP-1::GFP and suppression of the mutant *ham-1* Q.a reversal phenotype by *wsp-1* mutations demonstrate that the reversal of membrane extension contributes to the reversal of Q.a DCSA.

The normal Q.a myosin asymmetry in the *ham-1(gm279)* mutant came as a surprise for two reasons. First, our finding conflicts with the previous finding that NMY-2::GFP asymmetry is lost in the weak, partial loss-of-function *ham-1(cas46)* mutant [11]. Our analysis, by contrast, used a putative null *ham-1(gm279)* allele that lacks detectable HAM-1 protein [13, 14]. The difference in allele strength suggests that we should have observed a stronger myosin phenotype, but instead saw none. We have observed that the *nmy-2::gfp* transgene used in both studies can sometime produce lines that have reduced NMY-2::GFP fluorescence where we cannot detect asymmetry in Q.a. Perhaps a line with diminished fluorescence can account for the inability to detect the asymmetry in the *ham-1(cas46)* experiment. Alternatively, the *ham-1(cas46)* strain used by Feng et al. could contain a second mutation that altered NMY-2 localization or could result in neomorphic activity that alters NMY-2::GFP localization.

Second, observations in flies and worms suggest that asymmetric myosin locally inhibits membrane extension, restricting it to the opposite pole. In Drosophila neuroblasts, the myosin light chain localizes to the basal membrane during anaphase and is thought to inhibit basal cortical extension [27, 28]. The nonmuscle myosin NMY-2 localizes asymmetrically in Q.a but not Q.p, and Chromaphore Assisted Laser Inactivation (CALI) experiments indicates the anterior NMY-2 activity promotes Q.a DCSA, perhaps also by restricting membrane extension to the opposite pole [6].

Our finding demonstrates that NMY-2 localization at the anterior of a *ham-1* mutant Q.a is not sufficient to restrict membrane extension or WSP-1 localization. One possible explanation for this apparent paradox is that although present at the anterior of Q.a, NMY-2 is inactive. Asymmetric activation of NMY-2-mediated contraction by phosphorylation of myosin light-chain kinase, for example, could be defective in *ham-1* mutants. Another possibility is that other molecules that localize with NMY-2 regulate its activity and the localization of these molecules are disrupted in *ham-1* mutants. In Drosophila neuroblasts, the kinesin Pavarotti and Anillin colocalize asymmetrically with the Sqh [27, 28]. It is not known whether the *C. elegans* homologs of these proteins localize asymmetrically with NMY-2 in Q.a.

### HAM-1 suppresses Gα-mediated spindle movement in Q.a

Our findings support a hypothesis that a conserved mechanism for spindle movement leads to the posterior movement of the spindle in Q.p. In both the *C. elegans* zygote and Drosophila neuroblasts, Ga-GDP interacts with molecules that provide a physical bridge to dynein. This link between the membrane and microtubule asters provides the force that pulls the centrosome. The same Gα genes, *goa-1* and *gpa-16*, contribute to both zygote and Q.p DCSA. These Gαs bind directly to GPR-1/2, and the asymmetric activity of Ga-GDP in the zygote mediates the asymmetric distribution of mCherry::GPR-1 (reviewed in [2]). The asymmetric distribution of mCherry::GPR-1 in Q.p but not Q.a supports the hypothesis Gas mediate Q.p spindle asymmetry. Further support for this hypothesis comes from the suppression of the Q.a DCSA of *gpb-1* mutants by a *goa-1* mutation.

### The role of HAM-1 in DCSA

HAM-1 protein expression is dynamic. Regulation of HAM-1 levels could be transcriptional or posttranscriptional. Because the *ham-1* mutant Q.a phenotypes are similar to the wild-type Q.p phenotypes and because HAM-1 expression is restricted to Q.a, HAM-1 could be sufficient to generate a Q.a DCSA phenotype. Our attempts to address this problem have been difficult to interpret. Expression in the Q lineage of GFP-tagged HAM-1 made the Q.p division more symmetric, but untagged HAM-1 had no effect (data not shown). One possibility is that HAM-1 is regulated at the level of protein stability in Q.p and that the untagged HAM-1 is degraded, but the tagged versions of HAM-1 are more stable and available to alter the Q.p division. Alternatively, the tagged HAM-1 proteins could have neomorphic activity unrelated to their normal function.

The localization of HAM-1 is complex. Although antibodies only detect asymmetric location of endogenous HAM-1 to the posterior cortex of dividing cells [13, 14], GFP-tagged HAM-1 and cell fractionation studies show that HAM-1 also localizes to the nucleus [15]. Consistent with these findings, our full-length *Pham-1::GFP::ham-1* transgene, which retains partial function, localizes to the nucleus during interphase and to the cortex during mitosis. Mutating the two NLS sequences in a *ham-1 (Pegl-13::ham-1NLSmut)*transgene disrupts its ability to rescue the *ham-1* mutant Q.a DCSA defect. Similar to the Q.a requirement for HAM-1 nuclear localization, mutating the NLS sequences partially disrupted the ability of *ham-1* transgenes to rescue the mutant *ham-1* PLM defect [15]. Although this partial rescue could reflect HAM-1 functions outside of the nucleus, RNAi of the *C. elegans* exportin gene resulted in detectible nuclear GFP::HAM-1 when both NLS sequences wre mutated, suggesting that undetectable levels of this mutant HAM-1 could have been present in the nucleus and account for the residual HAM-1 function in the rescue experiment [15]. This possibility makes it impossible to know whether nuclear localization is absolutely or partially required for *ham-1* function in Q.a DCSA.

The presence of the Storkhead box, a winged-helix domain, suggests that HAM-1 is a transcription factor. Stox1, one of the two mammalian proteins containing a Storkhead box, has been shown to directly regulate transcription of target genes [33, 34]. Our finding that the transgene likely used in the ModEncode experiments to identify HAM-1 binding sites is a *mep-1::gfp* fusion reveals that there is currently no evidence for HAM-1 in transcription. It is noteworthy that winged-helix domain proteins can mediate other aspects of DNA regulation and can bind and regulate RNAs, and the winged-helix domain of some proteins do not bind nucleic acids but instead mediate protein-protein interactions [35].

Cortical localization requires two regions of HAM-1, sequences near the N-terminus defined by several missense mutations and a poly-proline motif defined by transgene deletions, have different requirements in Q.a DCSA and PLM development. The N-terminal mutations, in particular the *ham-1(n1811)* mutation, severely disrupts Q.a DCSA but has little effect on PLM development; by contrast, mutating the poly-proline motif in *ham-1* transgenes has a weak effect on rescue of the *ham-1* mutant Q.a DCSA defect but eliminates rescue of the *ham-1* PLM defect [15]. These findings suggest that although the N-terminal region and the poly-proline motif both promote cortical localization of HAM-1, the two sequences possess distinct functions.

Cortical and nuclear localization could reflect distinct HAM-1 functions that regulate DCSA. Alternatively, the two could be linked: HAM-1’s presence at one subcellular site could be required for its subsequent function at another site. For example, HAM-1 could be modified at the cortex so that it can act in the nucleus. Understanding how HAM-1 regulates DCSA will require identifying the molecules that interact with the Storkhead box, the N-terminus and the poly-proline motif.

## Materials and Methods

### Genetics

General handling and culture of nematodes were performed as previously described (Brenner 1974). The N2 Bristol line was used as wild type, and experiments were performed at 20°C unless otherwise noted. The following mutations and integrated arrays were used:

#### LG I

*gpa-16(it143)* (Bergmann et al, 2003, Johnston et al, 2008), *goa-1(n363)* (Segalat et al, 1995), *sptf-3(tm607)* [19]; *gmIs88[mab-5p::pig-1::gfp; pRF4(rol-6(su1006)); dpy-30p::NLS::DsRed2]*(Chien et al, 2013).

#### LG II

MosSCI transgenes integrated at the ttTi5605 site (see Table S1) (FrØkjÆr-Jensen *et al.* 2008), *gpb-1(pk44)* (Zwaal et al, 1996), *cdc-42(gk388)* [36].

#### LG III

*unc-119(ed3)* (Maduro and Pilgrim 1995), *rdvIs1* [*egl-17p::mCherry:his-24 + egl-17p::myristolated mCherry + pRF4*] (OU *et al.* 2010), *arx-7(ok1118)* ( *C. elegans* deletion mutant consortium).

#### LG IV

*pig-1(gm301*) (Cordes *et al.* 2006), *ham-1(cas27, cas46, cas137, gm214, gm267, gm279, n1438, n1811)* (Frank et al, 2005, Feng et al, 2013), *wsp-1(gm324, cas723[gfp::wsp‐1a knock‐in]) [21, 37]*.

#### LG V

*arx-2(ok1269)* [38], *zuIs45[nmy-2::gfp]* [39].

*LGX. gmIs81*[*mec-4p::mCherry*, *flp-12p::EBFP2*, *gcy-32p::gfp*, *egl-17p::gfp*] (Gurling *et al.* 2014), *nIs540[pig-1p::gfp; rol-4(su1006)]* [19].

Unmapped: *gmIs10[ham-1p::gfp::ham-1 + pRF4]* (this study), *ddIs79[mCherry::gpr-1 (synthetic, CAI 1.0, artificial introns) unc-119(+)]*[26].

### Neuron number scoring

The A/PVM, SDQR/L, A/PQR neurons were detected using the *gmIs81* reporter [16]. In *ham-1* mutants, extra neurons were found in close proximity to the normally unique AQR or PQR neurons.

### Plasmid construction and transgenics

The detail of DNA manipulations is presented in the supplemental material. Transgenic lines were generated by injecting plasmid DNA into the syncytial hermaphrodite gonad. MosSCI lines were generated as described by injecting the EG6699 *ttTi5605, unc-119(ed3)*mutants (FrØkjÆr-Jensen *et al.* 2014). All transgenes used in this study are described in Table S1.

### Neuroblasts daughters sizes measurement

Q.a and Q.p daughter cells sizes measurements were performed as previously described [10, 21, 40](Singhvi 2011, Teuliere 2014). For each genotype and ACD, the measured daughter cells size ratio was plotted on graphs where horizontal dotted lines indicate 1:1 ratios, and horizontal grey bars indicate the median of each ratio distribution.

### PIG-1::GFP transcriptional reporter quantification in Q cells

*pig-1* transcriptional activity measurements were performed using the*nIs540[pig-1p::gfp; rol-4(su1006)]*reporter introduced in wild-type and mutant strains also carrying the *rdvIs1* marker [19]. QR.a and p cells were observed after birth and before division as determined by their anterior position after migration and the division pattern of seam cells. GFP intensity was measured in the nucleus using ImageJ and was corrected for background fluorescence.

### Q neuroblast imaging

Q neuroblast imaging was performed as described [10, 16]. Time-lapse movies were either obtained by capturing single-plane images of dividing neuroblasts every 10 seconds to analyze membrane extension, chromosome movements, and NMY-2::GFP dynamics, or three planes Z-stacks (Z-step: 0.5 µm) every 20 seconds to image GFP::WSP-1 and GPR-1::mCherry fluorescence.

Image treatment and analysis were performed using ImageJ software. Movies were realigned using the StackReg plugin for ImageJ (http://bigwww.epfl.ch/thevenaz/stackreg/). Immobile cells labelled by the *rdvIs1* marker, such as V5L/R or QL.p, or by the *gmIs81* marker, such as ALMs or PLMs, were included in the field of acquisition as fiducial markers to confirm accurate realignment.

Plasma membrane extension was scored by subtracting binary images of the dividing cell outline from a metaphase image and a telophase image after time-lapse recording of the Q.a or Q.p division. Anterior (AE) and posterior (PE) extension areas represent the extension of the telophase membrane beyond the limits of the metaphase membrane.

Condensed chromatin in metaphase and anaphase is detectable by fluorescent labelling (*rdvIs1* marker) or fluorescence exclusion (*gmIs81* marker). To follow chromosome displacement in anaphase, a line was traced perpendicular to the metaphase plate from cell pole to cell pole, and kymographs were created using the Kymograph Plugin for ImageJ (http://www.embl.de/eamnet/html/body_kymograph.html). Chromosome displacement and time of displacement were measured from the center of the metaphase plate to the chromosome/midzone boundary when chromosomes have reached their final position at the end of anaphase.

NMY-2::GFP, GFP::WSP-1 and GPR-1::mCherry cortical and membrane intensities were measured in ImageJ on a single optical section using a 5 pixel-wide segmented line covering the anterior and posterior cortex of the dividing cell, and using the center of the metaphase plate and the spindle midzone, labelled by the *rdvIs1* or *gmIs81* transgenes as the anterior/posterior boundary. Chromosome separation was used as temporal reference to synchronise the movies.

GFP::HAM-1 expression pattern and subcellular localization were determined from the *gmIs10* transgene by collecting images at different stages of Q lineage development as determined by the morphology and position of the cells labelled by the *rdvIs1* transgene.

### RNA interference

RNAi was performed by feeding worms individual bacterial clones as described [41]. DCSA was analyzed in the L1 progeny of fed worms. The *goa-1* clone was from the library constructed in the Ahringer Lab [41, 42]. The *gpa-16* clone was a gift from Lesilee Rose. The negative control used in these experiments was a clone containing empty vector (L4440) in the bacterial host HT115 [41], the same host used for the RNAi feeding experiments. The *ham-1* clone was constructed by cloning a *ham-1* cDNA into the L4440 vector.

### Statistical analysis

Statistical analysis was performed using: the two-sample Z-test for proportions for extra neuron scoring; the Mann-Whitney U-test for cell size ratios, *pig-1p::GFP*, NMY-2::GFP, GFP::WSP-1 and GPR-1::mCherry intensity ratios; the Student T-test for chromosome displacement and membrane extension measurements.

### DNA manipulations and transgene construction

All DNA constructs and sequences used in this study are available upon request. PCR-amplification was performed using the Pfu Ultra high-fidelity polymerase (Invitrogen).

The GFP::HAM-1 fusion expressed from the *gmIs10* transgene was generated by PCR-engineering a PstI site at the amino terminus of *ham-1*, amplifying the sequence coding for GFP from the vector pPD95.02 (Addgene), and inserting the amplified GFP sequence into the engineered PstI site, placing GFP 7 amino acids downstream from the first methionine of HAM-1. The pKG17 plasmid contained the modified genomic fragment comprising the last 4 kb of *ham-1* 5’UTR, the *gfp::ham-1* fusion and the first 949 bp of *ham-1* 3’UTR. The pKG17 plasmid was coinjected at 50 ng/µl with the pRF4 plasmid at 100 ng/µl into N2 hermaphrodites gonads to generate an extra chromosomal array which was subsequently integrated into the genome by UV irradiation to create the *gmIs10* transgene.

The GFP::HAM-1 fusion used to express HAM-1 in Q.a cells was PCR-amplified from pNH144 (a kind gift from Dr. Nancy Hawkins) and consisted of the gfp cDNA directly fused to the 5’ end of the *ham-1* cDNA and Gateway cloned into pDONR221 (Invitrogen) to generate pENTR *gfp::ham-1*. Similarly, pENTR *Pmab-5* and pENTR *3’unc-54* entry clones were created by PCR-amplification of the 8.8 kb *mab-5* promoter and the *unc-54* 3’UTR, respectively, and Gateway cloning into pDONR P4-P1R and pDONR P2R-P3 (Invitrogen), respectively. A multisite 4-1-2-3 Gateway reaction was used to create *Pmab-5::gfp::ham-1* from these three entry clones and the destination vector pDEST R4R3 (Invitrogen).The *gmEx699* and *gmEx701* transgenes were created using standard procedures by coinjecting the *Pmab-5:gfp::ham-1* plasmid at 25ng/µl and the pCFJ90 *Pmyo-2::mcherry* plasmid (Addgene) at 3ng/µl into the gonads of *rdvIs1* hermaphrodites.

## Acknowledgements

Some nematode strains used in this work were provided by the Caenorhabditis Genetics Center, which is funded by the NIH Office of Research Infrastructure Programs (P40 OD010440). We thank Bob Horvitz and Guangshuo Ou for providing some of the strains used in this study. This work was supported by National Institutes of Health grant NS32057 to G.G.

## Supplementary Information

**Supplementary Figures and Legends**

**Fig S1.**
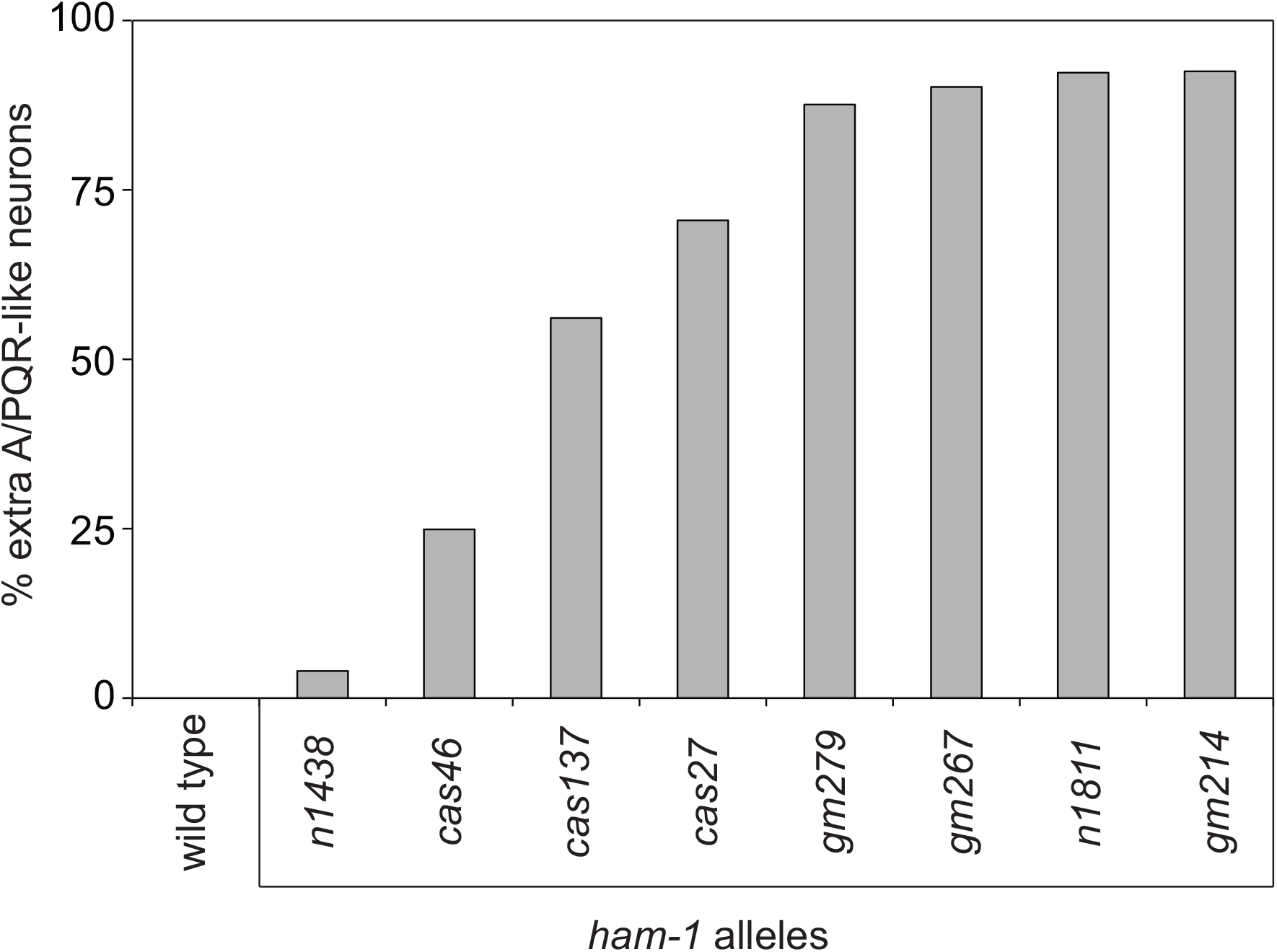
*ham-1* mutants have extra A/PQR-like neurons. Scoring of the extra A/PQR-like cells phenotype in wild-type and several *ham-1* mutants. The *ham-1(gm279)* null and strong hypomorphic *ham-1*(*n1811*, *gm267*, *gm214*) mutants had a high penetrance of extra A/PQR-like cells, indicating that the residues G47 (GD7D in *n1811*), G58 (G58R in *gm267*) and residues 379 to 389 (deleted in *gm214*) are essential for *ham-1* functions in the Q.a lineage [1]. One exception was the *ham-1(n1438)* mutation, a small deletion that removes promoter and 5’UTR sequences and that leads to an almost complete elimination of embryonic HAM-1 protein expression [1, 2]. Given that this mutation failed to induce penetrant A/PQR defects, it seems likely that these parts of the *ham-1* promoter and 5’UTR are not essential for HAM-1 expression in the Q lineage. Milder phenotypes were observed for G10 mutants (G10R in *cas27)* and (G10E in *cas137}*. Low penetrance defects were induced by a C-terminal truncation (*cas46)*, which removes the sequences for the C-terminus and 3’UTR (∆C33+3'UTR) as previously reported [3].

**Fig S2.**
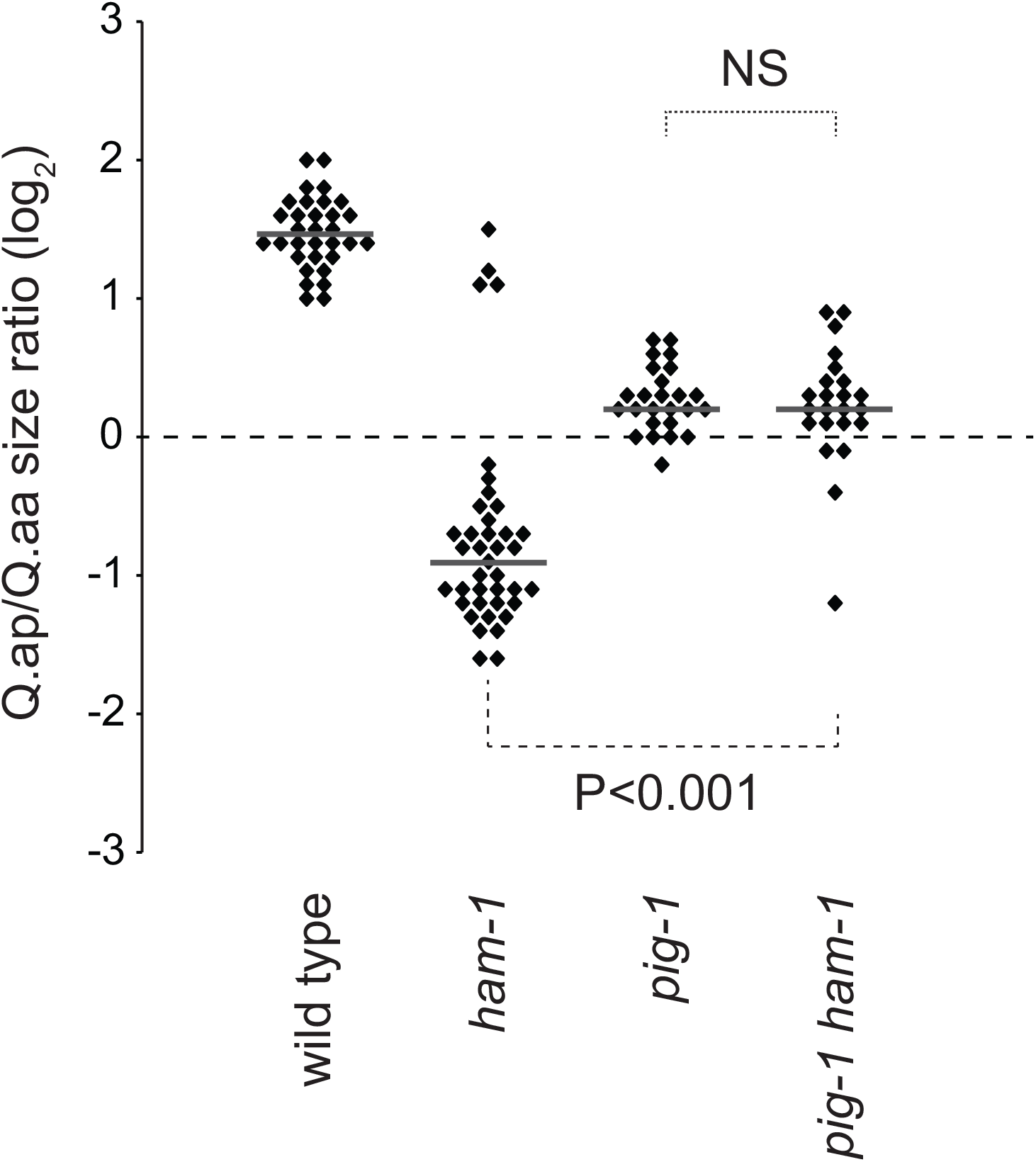
Genetic interaction between *ham-1* and *pig-1*. Distributions of Q.ap to Q.aa cell-size ratios in *ham-1*, *pig-1*, and *ham-1 pig-1* mutants carrying the *gmIs81[egl-17p::gfp]*marker. The *ham-1(gm279)* allele was used in these experiments. P-values obtained using the Mann-Whitney U-test are indicated. NS: not significant.

**Fig S3.**
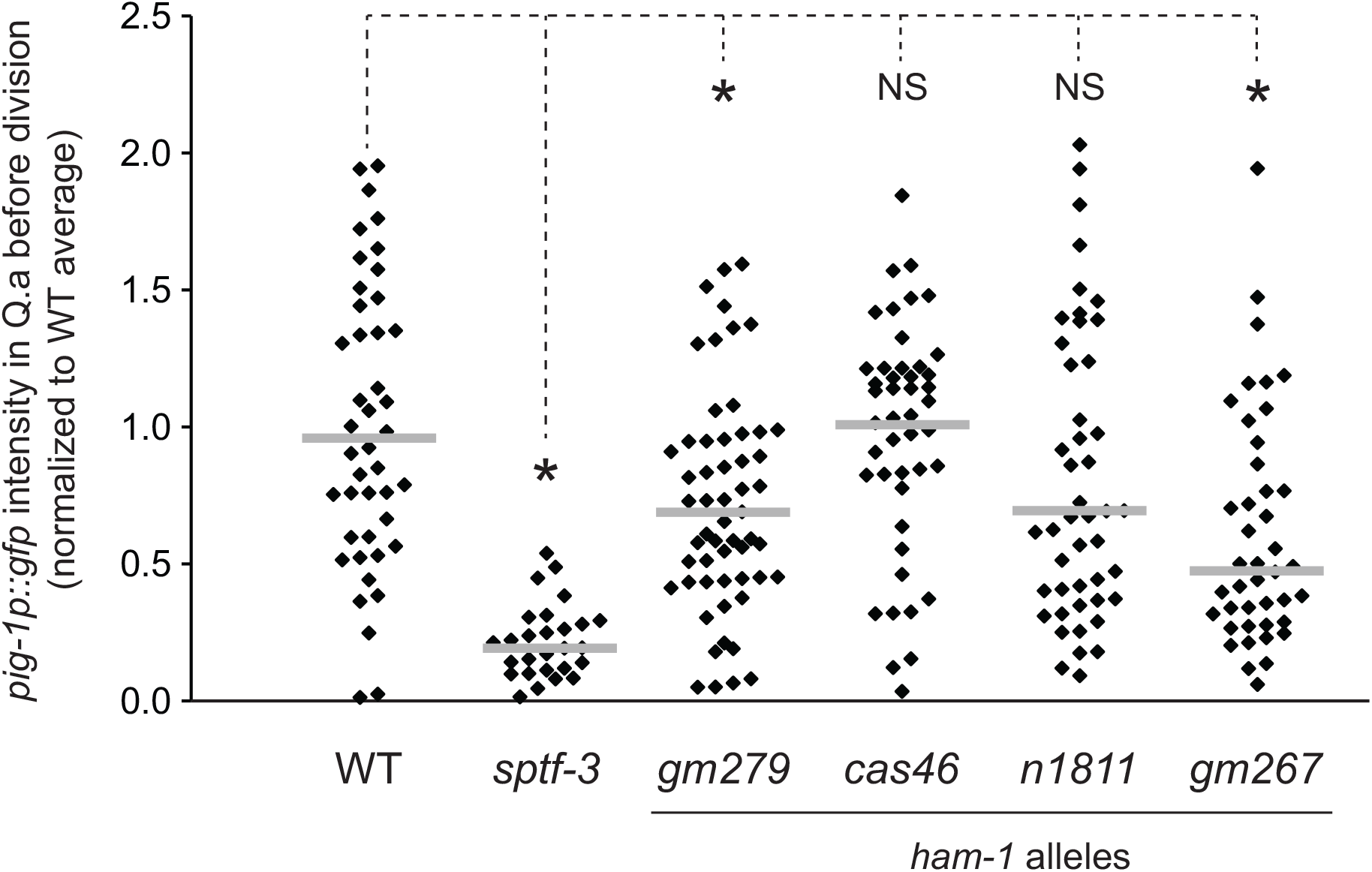
*pig-1* expression was not lost in *ham-1* mutants. *pig-1* transcriptional reporter *nIs540[pig-1p::gfp]* fluorescence intensity in QR.a before division. Each dot represents the intensity for one cell. The *sptf-3(tm607)* mutant was used as a positive control for loss of *pig-1* expression at the transcriptional level [4]. * P<0.01 using the Mann-Whitney U-test. NS: not significant.

**Fig S4.**
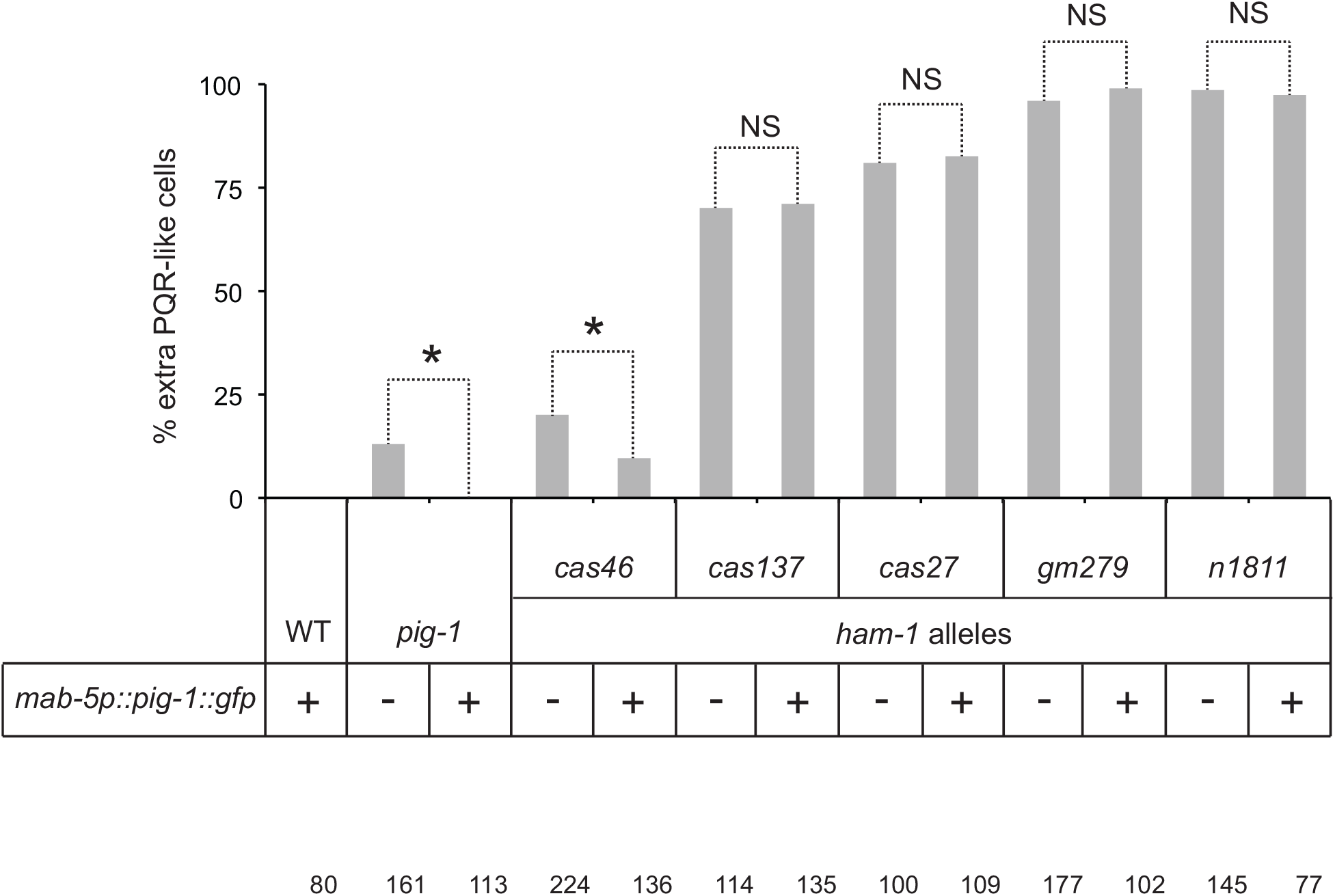
Transgenic PIG-1 expression in Q.a does not suppress the *ham-1* extra cell phenotype. Effect of PIG-1::GFP expression in the left Q lineage on the penetrance of extra PQR-like cells in *pig-1* and *ham-1* mutants. For each mutant, the extra-cell phenotype is compared in the absence (-) or presence (+) of the *gmIs88[mab-5p::pig-1::gfp]* transgene. The number of lineages scored is indicated under each bar. * P<0.001 using the Z-test for two proportions; NS: not significant.

**S5.**
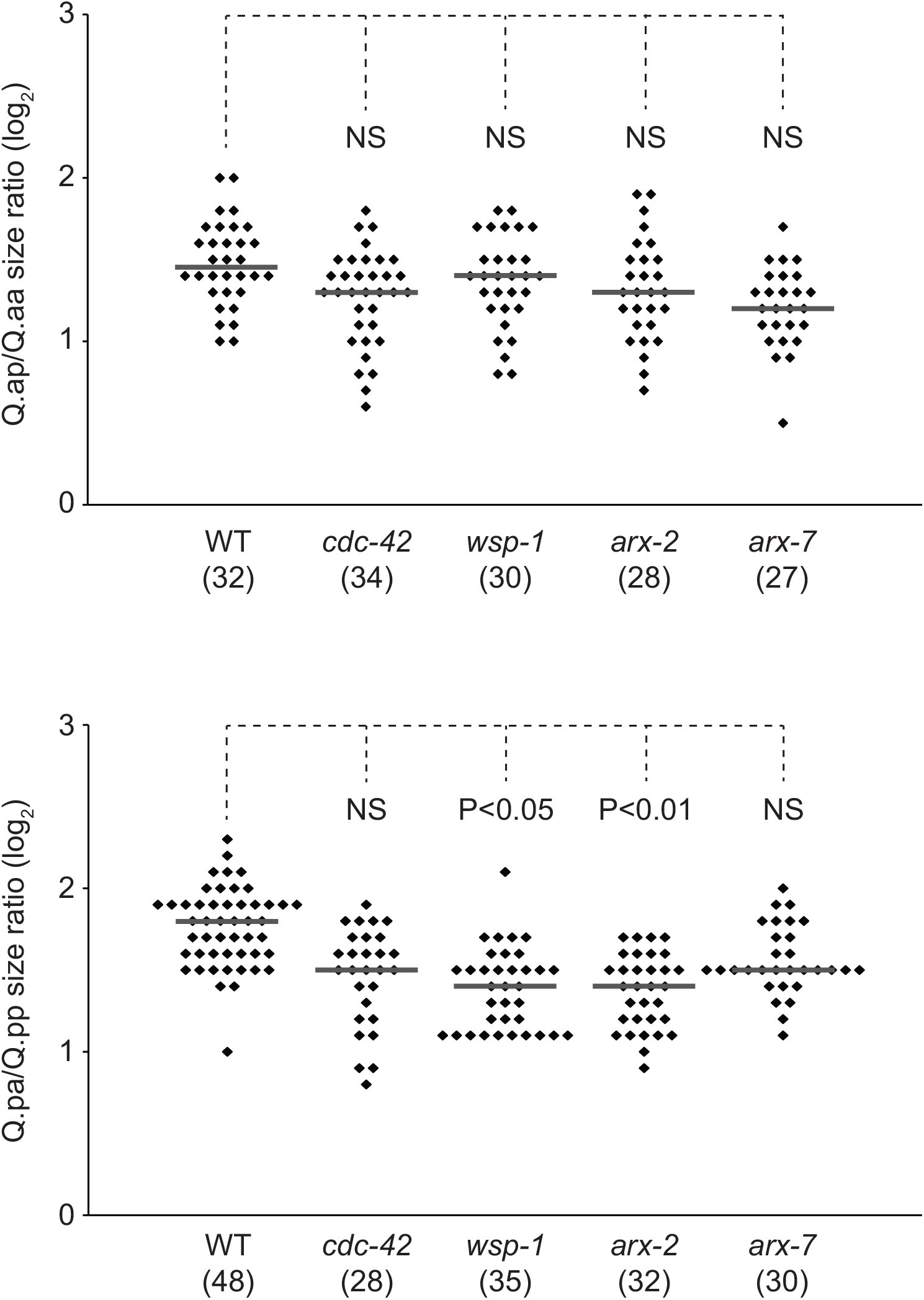
WSP-1 and the ARP-2/3 complex contribute to DCSA. (A, B) Distribution of cell-size ratios for Q.a (A) and Q.p (B) daughters. Cells were labelled using the *gmIs81[egl-17p::gfp]* marker. P-values obtained using the Mann-Whitney U-test are indicated. NS: not significant.

**Fig S6.**
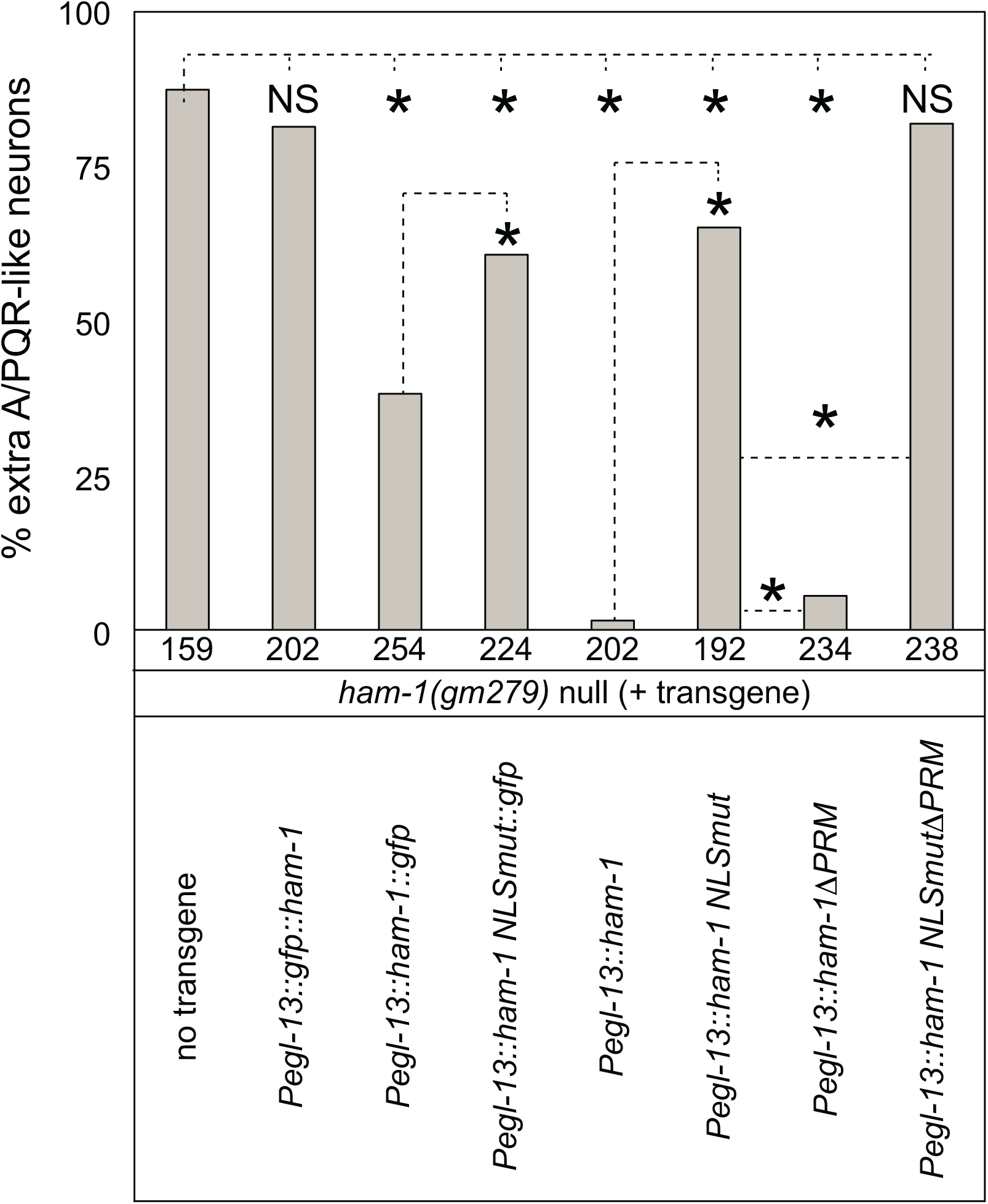
Transgenic HAM-1 expression in Q.a rescued the *ham-1* extra-cell phenotype. Effect of the expression of untagged wild-type and mutant HAM-1 in Q.a decreased on the penetrance of extra A/PQR-like cells in *ham-1*(*gm279*) mutants. The number of lineages scored is indicated under each bar. * P<0.001 using the Z-test for two proportions. NS: not significant.

**Fig S7.**
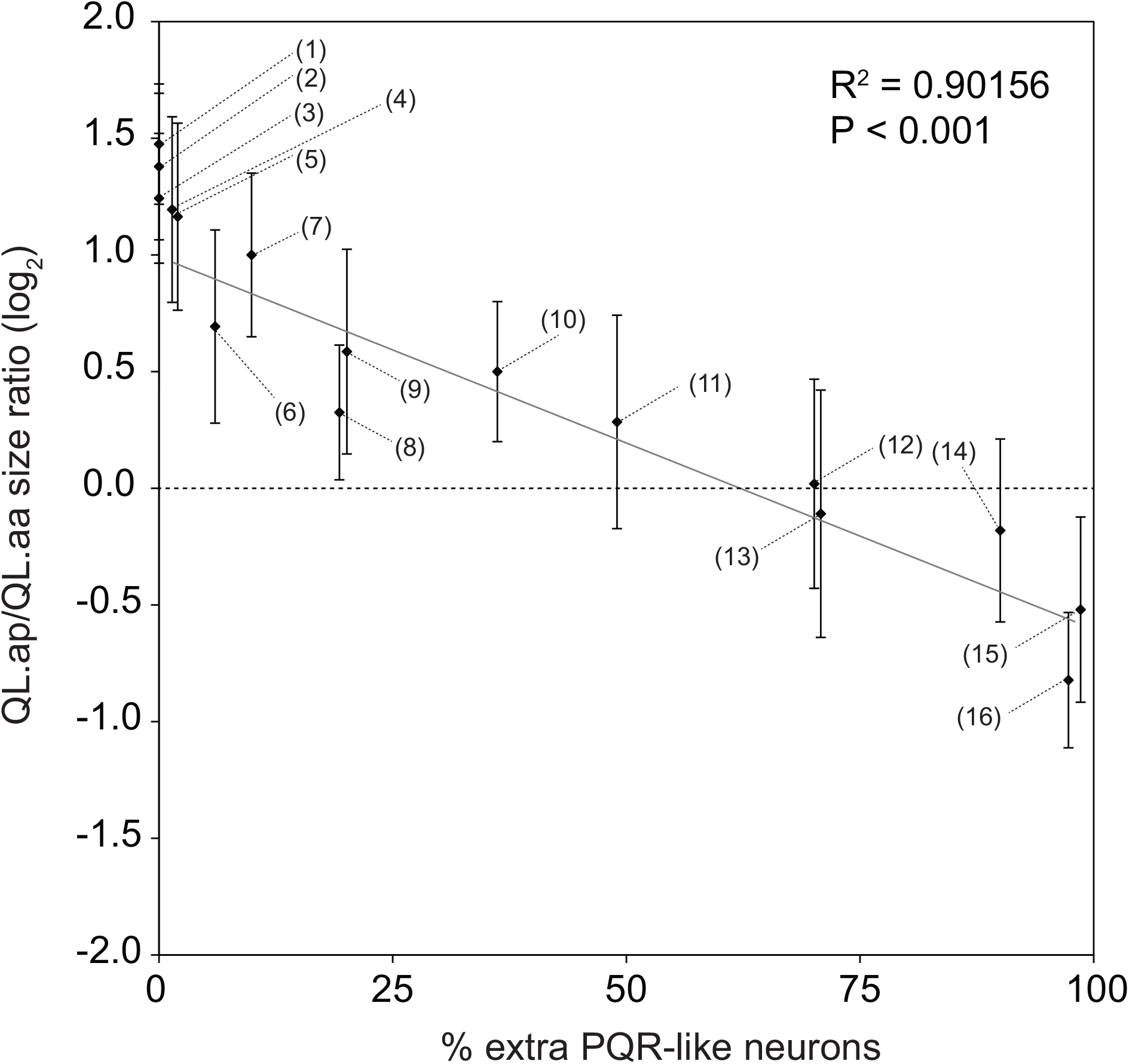
Correlation between Q.a DCSA and extra PQR-like neurons. The graph plots the Q.ap/Q.aa cell-size ratio (log2 representation) against the percent of Q.a lineages that produce extra surviving PQR-like neurons. Dots represent the mean ratio for each genotype indicated. Error bars indicate standard deviation. The linear regression (gray line) and the Pearson correlation coefficient (R) and P-value (P) are indicated on the graph. Genotypes: (1) wild type, (2) *gmSi51; ham-1(cas46)*, (3) *gmSi53; ham-1(cas46)*, (4)*gmSi51; ham-1(n1811)*, (5)*gmSi51; ham-1(gm279)*, (6) *gmSi151; ham-1(gm279)*, (7)*ham-1(gm413)*, (8)*pig-1(gm301)*, (9)*ham-1(cas46)*, (10)*gmSi140; ham-1(gm279)*, (11)*ham-1(gm279); gmIs10*, (12)*ham-1(cas137)*, (13)*gmSi53; ham-1(gm279)*, (14)*gmSi153; ham-1(gm279)*, (15)*ham-1(n1811)*, (16)*ham-1(gm279)*.

